# Non-canonical immune response to the inhibition of DNA methylation via stabilization of endogenous retrovirus dsRNAs

**DOI:** 10.1101/2020.06.21.163857

**Authors:** Yongsuk Ku, Joo-Hwan Park, Ryeongeun Cho, Yongki Lee, Hyoung-Min Park, MinA Kim, Kyunghoon Hur, Soo Young Byun, Jun Liu, David Shum, Dong-Yeop Shin, Youngil Koh, Je-Yoel Cho, Sung-Soo Yoon, Junshik Hong, Yoosik Kim

**Affiliations:** Department of Chemical and Biomolecular Engineering and KAIST Institute for Health Science and Technology (KIHST), Korea Advanced Institute of Science and Technology (KAIST), Daejeon, Korea; Department of Internal Medicine, Seoul National University College of Medicine, Seoul National University Hospital, Seoul, Korea; Cancer Research Institute, Seoul National University Hospital, Seoul, Korea; Department of Biochemistry, BK21 Plus and Research Institute for Veterinary Science, School of Veterinary Medicine, Seoul National University, Seoul, Korea; Screening Discovery Platform, Translation Research Division, Institut Pasteur Korea, Seongnam, Gyeonggi, Korea

**Keywords:** Endogenous retroviruses, DNA-methyltransferase inhibitor, dsRNA-binding protein, non-coding RNA, innate immune response, acute myeloid leukemia, myelodysplastic syndromes, Staufen, TINCR

## Abstract

5-Aza-2′-deoxycytidine, also known as decitabine, is a DNA-methyltransferase inhibitor (DNMTi) used to treat acute myeloid leukemia (AML). Decitabine activates the transcription of endogenous retroviruses (ERV), which can induce an immune response by acting as cellular double-stranded RNAs (dsRNAs). Here, we employ an image-based screening platform to identify dsRNA-binding factors that mediate the downstream effect of ERV induction. We find that Staufen1 (Stau1) knockdown decreases the interferon signature and rescues decitabine-mediated cell death. Moreover, Stau1 directly binds to ERV RNAs and stabilizes them, together with a long non-coding RNA TINCR. We further show that TINCR enhances the interaction between Stau1 and ERV RNAs. Analysis of a clinical patient cohort reveals that AML patients with low Stau1 and TINCR expressions exhibits inferior treatment outcomes to the DNMTi therapy. Our study reveals that decitabine-mediated cell death is a consequence of complex interactions among different dsRNA-binding proteins for access to their common dsRNA targets.

**HIGHLIHGTS:**

- Image-based RNAi screening reveals multiple dsRBPs regulate response to decitabine
- Stau1 binds to ERV RNAs and affects their stability and subcellular localization
- TINCR binds to Stau1 and enhances Stau1-ERV interactions
- AML/MDS patients with low Stau1 and TINCR expressions show poor response to DNMTis

## INTRODUCTION

Long double-stranded RNAs (dsRNAs) are duplex RNAs that can trigger an immune response by serving as an antigen for pattern recognition receptors (PRRs) (Schlee and Hartmann, 2016). These RNAs are strongly associated with viral infection because they are produced during bidirectional transcription of viral genome for DNA viruses and as replication byproduct for RNA viruses, especially for positive-sense single-stranded RNA viruses(Pettersson and Philipson, 1974; Weber et al., 2006). The human genome encodes four different PRRs that can recognize long viral dsRNAs. They include melanoma differentiation-associated protein 5 (MDA5), protein kinase R (PKR), retinoic acid-inducible gene I (RIG-I), and toll-like receptor 3 (TLR3) (Akira et al., 2006; Hur, 2019). Although both DNA and dsRNA form helical structures, dsRNA exists as an A-form helix with a deep and narrow major groove where proteins have limited access to the individual bases (Masliah et al., 2013; Vukovic et al., 2014). Consequently, these dsRNA-sensors recognize the structural features of the target dsRNAs rather than their specific nucleotide. For example, PKR binds to dsRNAs longer than 33 base-pairs (bp), which leads to dimerization and autophosphorylation of the protein (Kostura and Mathews, 1989; Lemaire et al., 2008; Thomis and Samuel, 1993). When phosphorylated, PKR becomes active and induces phosphorylation of a number of downstream substrates to suppress global translation and to initiate an antiviral response (Wek et al., 2006). Similarly, MDA5 oligomerizes on dsRNAs longer than 100 bp, which then signals through mitochondrial antiviral signaling proteins (MAVS) to induce type I interferon production (Wu et al., 2013). This structure-based interaction between dsRNAs and PRRs may allow these immune response proteins to respond to a wide range of viruses with different genomic sequences.

Historically, long dsRNAs were believed to be a signature of viral infection. However, recent evidence suggests that the human genome contains numerous repeat elements that, when transcribed, can adopt a double-stranded secondary structure (Athanasiadis et al., 2004; Lander et al., 2001; Liu et al., 2019). Moreover, these cellular dsRNAs are also recognized by dsRNA sensors of the innate immune response proteins and activate them in uninfected cells. The most well-characterized cellular dsRNAs are from small-interspersed nuclear elements (SINEs), which are mostly Alus in primates. Alus occupy more than 10% of the human genome and are located primarily on the non-coding regions of the coding genes such as introns and untranslated regions (UTRs) (Lander et al., 2001; Rubin et al., 1980). When transcribed with the host RNA, two Alu elements with the opposite orientation can form intra- or intermolecular dsRNAs that are recognized by PKR and MDA5 as well as by other dsRNA-binding proteins (dsRBPs) such as adenosine deaminase acting on RNA (ADAR), which disrupts dsRNA structure by converting adenosine to inosine, and Staufen (Stau), which affects Alu dsRNA localization and stability (Ahmad et al., 2018; Bahn et al., 2015; Chen and Carmichael, 2008; Elbarbary et al., 2013; Kim et al., 2014; Liddicoat et al., 2015). In addition to Alus, recent studies have shown that RNAs from long-interspersed nuclear elements (LINE), endogenous retroviruses (ERVs), and even mitochondrial RNAs (mtRNAs) can act as cellular dsRNAs to regulate antiviral signaling under physiological conditions (Chiappinelli et al., 2015; Dhir et al., 2018; Kim et al., 2018; Roulois et al., 2015). More importantly, these cellular dsRNAs are strongly associated with human diseases where the misactivation of PKR and MDA5 are reported (Kim et al., 2019). For example, mutation in ADAR and subsequent immune response by unedited Alu dsRNAs is associated with Aicardi-Goutières syndrome while patients with mutations in PNPase, which removes mtRNAs, show strong interferon signature (Ahmad et al., 2018; Dhir et al., 2018; Rice et al., 2012).

Since these cellular dsRNAs can activate immune response proteins, the expression of these RNAs is usually silenced through epigenetic regulation. It has been shown that treating colorectal cancer cells with a DNA-methyltransferase inhibitor (DNMTi), decitabine or azacitidine, induces the expression of ERV RNAs, leading to MDA5 activation and subsequent cell death (Chiappinelli et al., 2015; Roulois et al., 2015). In addition, activation of the immune response system through dsRNA induction also resulted in increased susceptibility of cancer cells to immunotherapy, suggesting the importance and clinical potential of the dsRNA regulation. Yet, our understanding of ERV RNAs at the post-transcriptional level is still limited. Considering that many dsRBPs, such as ADAR and Stau, share a similar target pool as that of MDA5 and PKR, these proteins may also bind to ERV RNAs and affect cellular response to the DNMTi therapy. In addition, DNMTis are being used to treat myelodysplastic syndrome (MDS) and acute myeloid leukemia (AML) patients. Therefore, investigation on the regulatory factors of ERV dsRNAs may lead to the development of diagnostic markers to predict the responsiveness to the DNMTi therapy.

In this study, we employ an image-based RNA interference (RNAi) screening platform to identify dsRBPs that affect immune response by cellular dsRNAs. We analyze the effect of the knockdown of nine dsRBPs individually as well as in pairs to examine their synergistic or antagonistic regulation in response to the DNMTi treatment. We find that Stau1 knockdown significantly rescues cell death and immune response by decitabine treatment. We further show that Stau1 directly binds to ERV dsRNAs and regulates their stability and subcellular localization. Using formaldehyde-mediated crosslinking precipitation and sequencing (fCLIP-seq) of a dsRNA recognizing antibody, we find that Stau1-mediated stabilization of ERV RNAs and subsequent immune response occur in a genome-wide manner. Furthermore, we show that Stau1 stabilizes ERV RNAs through a cofactor TINCR non-coding RNA, which forms a complex with Stau1 and enhances the interaction between Stau1 and ERV RNAs. Lastly, we provide a clinical significance of our findings by analyzing a clinical cohort of AML and MDS patients who received the DNMTi therapy. We find that patients with lower Stau1 and TINCR expression showed significantly inferior treatment outcomes, both in progression-free survival (PFS) and overall survival (OS) rates. Collectively, our study establishes the functional role of Stau1 in its stabilization of cellular dsRNAs and provides the significance of dsRNA regulation in maintaining proper cellular physiology.

## RESULT

### Multiple dsRBPs affect response to low-dose exposure to decitabine

Previous studies have established that transient low-dose treatment of a DNMTi decitabine to colorectal cancer cells results in cell death via viral mimicry (Chiappinelli et al., 2015; Roulois et al., 2015). Here, we applied a similar experimental system where we treated HCT116 cells with 0.5 μM of decitabine for 24 h and cultured cells in a fresh media for up to 4 days (total of 5 days). As shown by previous studies, we also observed delay in doubling time starting 2 days after the treatment and significant cell death by the 5^th^ day (Fig. 1A). We extracted total RNAs from decitabine treated cells and confirmed that many ERV RNAs were strongly induced, which resulted in the increased expression of several interferon-stimulated genes (ISGs) (Fig. 1B and 1C). Indeed, in MDA5-deficient HCT116 cells, the induction of ISGs by decitabine was nearly abrogated, which resulted in increased cell proliferation (Fig. 1C and 1D). However, the degree of the increase in cell proliferation in MDA5-deficient cells was too small. This indicates that there may exist other factors that regulate response to DNMTis.

**Figure 1.**
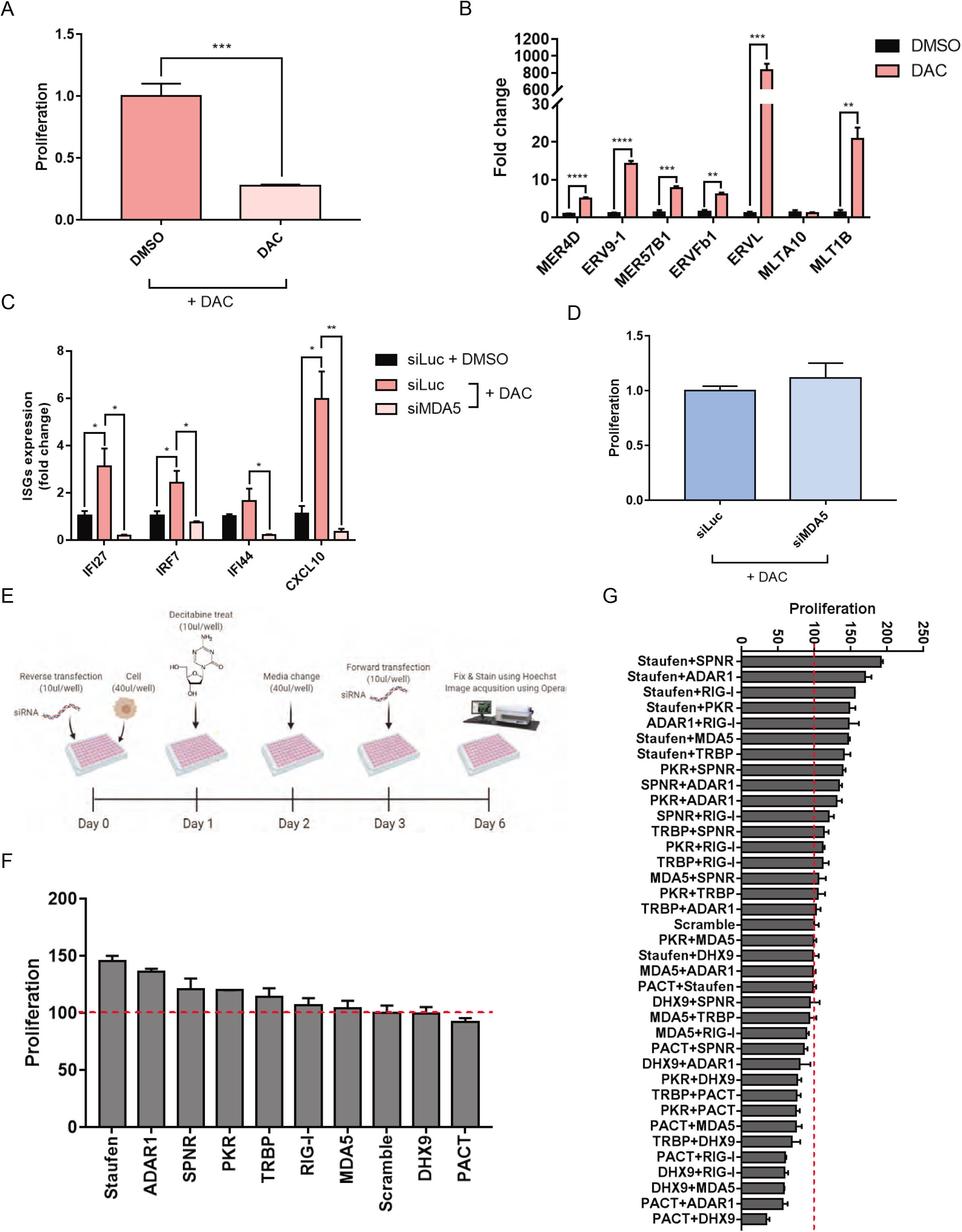
Several dsRBPs affect cell death by transient low dose exposure to decitabine. (A) Transient low dose exposure to decitabine for 5 days resulted in a significant decrease in proliferation measured by SRB assay. (B) The induction of several ERV RNAs by decitabine treatment was confirmed using qRT-PCR. (C) Moreover, increased ERV RNA levels lead to MDA5-dependent induction of ISGs. (D) The knockdown of MDA5 partially rescued cell death by decitabine. (E) A schematic of the image-based dsRBP screening employed in this study. (F, G) Cell proliferation results of image-based RNAi screening when dsRBPs were knocked down individually (F) or in pairs (G). In all experiments, an average of 3 biological replicates are shown with error bars denote s.e.m. A student’s t-test was performed for statistical analysis. *p-value < 0.05; **p-value < 0.01 ***p-value < 0.001 ****p-value < 0.0001

One strong candidate was PKR, another member of the innate immune response system that interacts with long dsRNAs. Using PKR fCLIP-seq, we have previously shown that PKR can also bind to ERV RNAs (Kim et al., 2018). In addition, considering that PKR only requires dsRNAs longer than 33 bp to become activated (Patel et al., 1995), the PKR signaling system is likely to be regulated by ERV RNAs during decitabine treatment (Fig. S1A). Indeed, we found that phosphorylated PKR (pPKR) as well as phosphorylation of eIF2α, a main downstream substrate of PKR, were increased 5 days after decitabine treatment (Fig. S1B and S1C). Moreover, PKR-deficient cells show significant rescue in cell viability even stronger than that of MDA5 knockdown (Fig. S1D-S1F). Therefore, both PKR and MDA5 are involved in cellular response to the DNMTi.

The human genome encodes multiple dsRBPs that share a similar dsRNA binding domain (dsRBD) as that of PKR. Previous biochemical studies have established that these dsRBDs can only recognize the structural features rather than specific RNA sequences (Saunders and Barber, 2003). Consistent with this, previous high-throughput studies have shown that dsRBPs, such as adenosine deaminase acting on RNA (ADAR), Staufen (Stau), DHX9, and PKR, all bind and regulate or regulated by dsRNAs formed by Alu repeats (Aktas et al., 2017; Bahn et al., 2015; Kim et al., 2018; Ricci et al., 2014). Considering that PKR could bind to ERV RNAs and the downregulation of PKR significantly rescued cell death by decitabine treatment, we hypothesized that many other dsRBPs would participate in the response to ERV induction by decitabine treatment. Moreover, these dsRBPs may work together or against each other in response to increased ERV RNA expressions.

To investigate and identify dsRBP factors that regulate decitabine-mediated cell death, we employed an image-based high-throughput screening platform. We first performed an initial screening of 14 dsRBPs and chose nine based on their known function and their ability to mediate the downstream effect of decitabine. We used the same experimental set up used for our analysis of MDA5 and PKR, where we transfected cells with an siRNA targeting a specific dsRBP twice over a 6-day period in order to ensure suppression of the target protein expression throughout the experiment (Fig. 1E). We found that Stau1 and ADAR1 knockdown resulted in the strongest degree of rescue in cell viability while no dsRBP alone can enhance the apoptotic effect of decitabine (Fig. 1F). Consistent with our earlier experiments, MDA5 knockdown only had a moderate effect while PKR knockdown had an intermediate effect; it was stronger than MDA5, but weaker than that of Stau1 or ADAR knockdown. When we performed the same screening assay using a combinatorial knockdown of two dsRBPs, we could clearly observe synergistic effects. Most of the dsRBPs worked together with Stau1 in that when Stau1 and another dsRBP were both knocked down, a significant rescue was observed in cell death (Fig. 1G). Interestingly, the combinatorial knockdown of DHX9 and protein activator of PKR (PACT) significantly enhanced the effect of decitabine. In this context, DHX9 and PACT were working together to sensitize cells toward decitabine. Antagonistic effects were also observed where the knockdown of PACT or DHX9 cancelled the rescue effect of Stau1 knockdown (Fig. 1G).

### Stau1 regulates the expression level and subcellular localization of ERV RNAs

Our image-based dsRBP screening revealed that Stau1 knockdown resulted in the largest degree of the rescue from decitabine-mediated cell death. We focused on Stau1-mediated regulation and asked how Stau1 affects the cellular response to decitabine, especially with respect to the induction of ERV RNAs. We designed multiple siRNAs that target either CDS or 3’ UTR of the Stau1 mRNA and confirmed their knockdown efficiencies through western blotting (Fig. 2A). We then confirmed that Stau1 knockdown almost completely rescued cell death by decitabine treatment, which is consistent with our image-based screening result (Fig. 2B). More importantly, we showed that Stau1 knockdown alone without decitabine did not affect the cell proliferation rate, indicating that the observed effect is not through Stau1 knockdown affecting the cell doubling time independent of decitabine. To investigate the mechanism, we first asked whether Stau1 could directly interact with ERV RNAs. We performed Stau1 fCLIP followed by qPCR analysis and showed that Stau1 strongly interacted with several ERV RNAs, particularly when decitabine was treated (Fig. 2C). Next, we examined the expression of ERV RNAs in Stau1-deficient cells. To our surprise, we found that, when Stau1 expression was reduced, the level of most ERV RNAs was significantly decreased (Fig. 2D). This result was rather surprising because Stau1 is known to destabilize dsRNAs by recruiting UPF1 and triggering Staufen-mediated decay (SMD) (Gong and Maquat, 2011). On the contrary, our results show that, in the case of ERV RNAs, Stau1 stabilizes the dsRNA expression.

**Figure 2.**
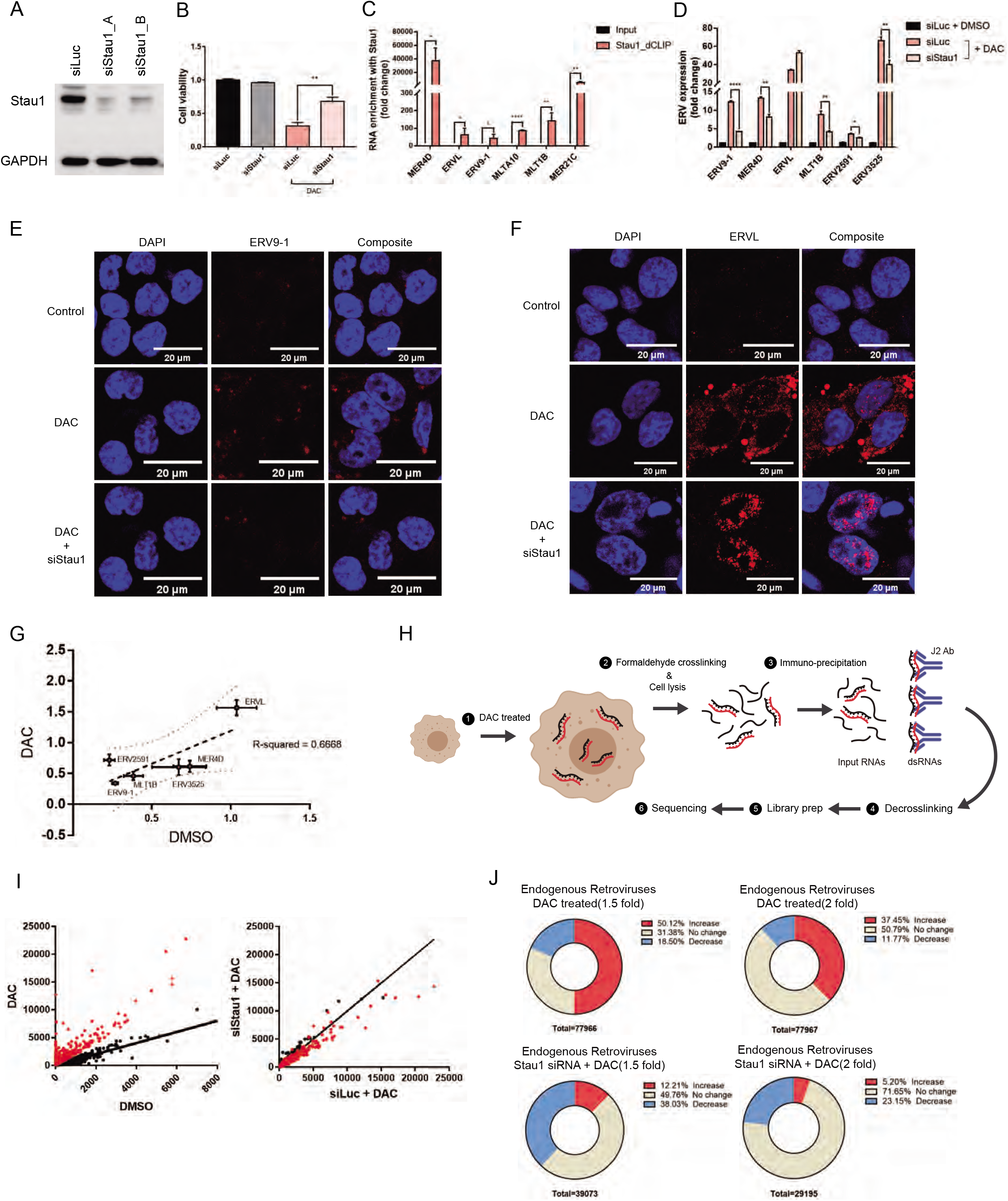
Stau1 interacts with ERVs and regulates their expression and subcellular localization. (A) Western blot result to confirm the knockdown efficiency of siRNAs used in this study. siStau1_A targets the 3’ UTR of Stau1 mRNA while siStau1_B targets the coding region of the mRNA. We present the data using siStau1_A for the rest of the study, but siStau1_B also yielded similar results. (B) The knockdown of Stau1 significantly rescued cell death from decitabine treatment. *n*= 3 and error bars denote s.e.m. (C) Direct interaction between Stau1 and ERV RNAs was confirmed using RNA-IP after formaldehyde crosslinking. *n*= 4 and error bars denote s.e.m. (D) Knockdown of Stau1 resulted in decreased expression of most ERV RNAs examined. *n*= 3 and error bars denote s.e.m. (E, F) RNA FISH of ERV9-1 (E) and ERV-L (F) revealed that Stau1 could affect expression and subcellular localization of ERV RNAs. (G) Stau1 knockdown also affected the basal expression of ERV RNAs without decitabine treatment. Fractional change of ERV RNA expression levels assessed by qRT-PCR for DMSO control and decitabine treated samples is shown. The two conditions show a good positive correlation with R^2^ = 0.6668. (H) A schematic for the J2 fCLIP-seq process. (I) Distribution of ERV RNA expression in the J2 fCLIP-seq sequencing result. Sequencing results were normalized with the GAPDH expression level. (J) Pie chart of endogenous ERV RNA expression change with decitabine and in Stau1 knockdown cells.

For ERV9-1, we designed RNA probes and performed fluorescent in-situ hybridization (FISH) to visualize the expression pattern of the RNA. While the control cells did not show any ERV9-1 RNA FISH signal, decitabine treated cells show strong red fluorescent signals throughout the cytosol, indicating the induction of ERV9-1 (Fig. 2E). However, in Stau1 knockdown cells, the ERV9-1 RNA FISH signal was almost completely eliminated (Fig. 2E). Furthermore, using RNA-FISH, we also examined the expression of ERV-L RNA because ERV-L was the only type of ERV element whose expression was not decreased in Stau1-deficient cells. Surprisingly, we found that when Stau1 was knocked down, the subcellular localization of ERV-L RNA was affected where cytosolic ERV-L RNAs were sequestered inside the nucleus (Fig. 2F). In this context, Stau1 may be important in the cytosolic export of ERV-L RNA, similar to its role in the regulation of dsRNAs formed by inverted Alu repeats (Elbarbary et al., 2013). Lastly, we examined whether Stau1 could affect the basal expression of ERV RNAs without decitabine induction. We found that for most ERV RNAs examined, Stau1 knockdown decreased their basal expression levels. When we plotted the percent change of ERV RNAs in DMSO control versus decitabine treated cells, the two conditions show a good positive correlation, indicating that Stau1 might be a general regulator of ERV dsRNA expression (Fig. 2G).

Encouraged by these results, we performed a genome-wide analysis to examine and identify ERV elements that are regulated by decitabine and by Stau1. The human genome encodes about 98,000 ERV elements, but it is unclear the percent of these ERV elements potentially induced by DNMTi treatment and the fraction of these ERV RNAs that are also regulated by Stau1. First, we analyzed ERV RNAs using total RNA-seq. However, the sequencing reads mapped to the ERV elements were too few that we could not obtain statistically meaningful results for most ERVs. Instead, we enriched dsRNAs through J2 fCLIP and analyzed the enriched dsRNAs by high-throughput sequencing (Fig. 2H). Through this approach, we were able to map a significant number of sequencing reads to about 78,000 ERV elements. Sequencing read accumulation and change on an exemplary ERV element (ERV1_LTR1_3525) are shown in Figure S2A. Structural prediction using RNAFold clearly predicts that this ERV element is likely to generate the dsRNA structure when transcribed (Fig. S2B). Among the 78,000 identified ERV elements, we found that about 50% (~39,000) ERV elements showed at least a 1.5-fold increase in RNA expression (Fig. 2I and 2J). Out of these ERV elements, about 40% showed at least a 1.5-fold decrease in RNA expression when Stau1 was knocked down. Similar conclusions could be reached even when we increased the threshold to 2-fold induction; about 37.5% of mapped ERV elements showed an increase in RNA expression upon decitabine treatment and out of these, 23% showed at least a 2-fold decrease in expression when Stau1 was knocked down (Fig. 2I and 2J). Of note, when we analyzed our sequencing data for other retrotransposon elements that can generate dsRNAs, such as SINE and LINE, we found that the proportion of ERV was decreased in decitabine treated cells (Fig. S2C). Moreover, the proportion of individual retrotransposable elements was restored when Stau1 was knocked down. Although the current study is focused on ERV RNAs, our sequencing result indicates that decitabine may regulate other cellular dsRNAs, particularly those from retrotransposable elements and that Stau1 also affects the expression level of cellular dsRNAs in addition to ERVs.

### Restoring Stau1 can rescue the ERV expression

As our analysis is based on RNAi-mediated gene knockdown, we further complemented our data by generating Stau1 knockout (KO) HCT116 cells using the CRISPR/Cas9 system. We designed an sgRNA targeting the exon 7 of Stau1 locus and successfully generated a frameshift mutant cell line (Fig. S3). The western blotting analysis showed the complete elimination of the band corresponding to Stau1 protein (Fig. 3A). Of note, we were able to detect a weak band slightly smaller than the size of wild-type Stau1. We further confirmed whether this residual band was from truncated Stau1 by transfecting cells with an siRNA that targets the 3’ UTR of Stau1 mRNA. There was no change in the band intensity, indicating that the observed band was likely a background signal (Fig. 3A). In Stau1 KO cells, we examined the induction of ERV RNAs upon transient exposure to low-dose decitabine. We found that with the exception of ERV-L, the magnitude of ERV induction was significantly reduced in the KO cells (Fig. 3B). This is consistent with the knockdown result that Stau1 deficient cells showed decreased expression of ERVs.

**Figure 3.**
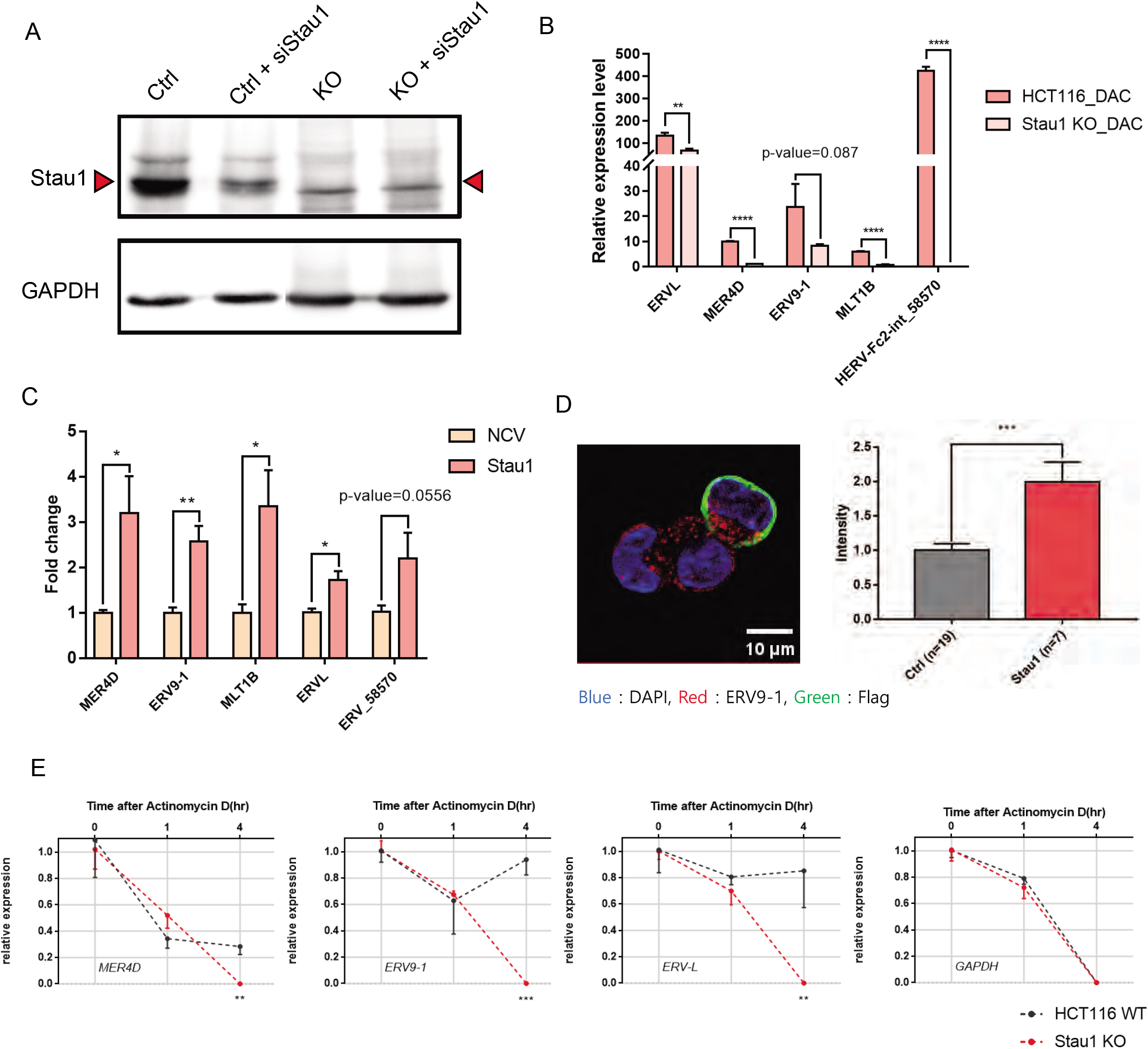
Analysis of Stau1 KO cells reveals the stabilization of ERV RNAs by Stau1. (A) Stau1 KO HCT116 cell is generated by targeting the exon 7 of Stau1 using CRISPR. Western blot analysis shows the elimination of Stau1 band in the lysates from the KO cells. (B) Compared to the parental cells, Stau1 KO cells showed decreased levels of ERV RNAs when treated with decitabine. *n= 3* and error bars denote s.e.m. (C) Overexpression of Stau1 in the KO cells resulted in increased expression of all ERVs examined. NCV indicates the transfection of the blank vector. (D) Increased expression of ERV9-1 RNA was also shown through RNA-FISH image analysis. Stau1 expressing cells were identified by staining cells with the Flag antibody, shown in green. Quantified fluorescent signals are shown in the right. *n*= 19 (Ctrl), *n*= 7 (Stau1) and error bars denote s.e.m. (E) ERV RNA expression changes after treating parental and KO cells with ActD. *n= 3* and error bars denote s.e.m. *p-value < 0.05; **p-value < 0.01 ***p-value < 0.001 ****p-value < 0.0001

We then performed rescue experiments by overexpressing wild-type Stau1 in the KO cells. We found that for all of the ERVs examined, overexpressing wild-type Stau1 resulted in increased expression (Fig. 3C). For ERV9-1, increased RNA expression was further confirmed through RNA-FISH. For this analysis, we identified Stau1 expressing cells using green fluorescent signals by targeting the Flag tag on the exogenous Stau1 and compared the red fluorescent signal intensities with those from the cells that do not express Stau1 protein. We found that the Stau1 expressing cells showed significantly higher red fluorescent intensity, indicating the higher expression of ERV9-1 RNAs (Fig. 3D).

We also utilized the KO cell to further examine how Stau1 knockdown decreased the expression of ERV RNAs. We treated cells with actinomycin D (ActD) to shut down the transcription and analyzed the stability of ERV RNAs over time. We examined ERV RNA expression normalized to that of the U6 small nucleolar RNA, which is transcribed by polymerase III and is not affected by ActD. Compared to the parental cells, the KO cells showed decreased expression of ERV RNAs upon ActD treatment, indicating faster degradation of the RNAs (Fig. 3E). Therefore, Stau1 affects the ERV RNA expression by directly binding and stabilizing the transcript.

### Stau1 affects the immune response by decitabine

Our results indicate that Stau1 knockdown decreases ERV expressions in a genome-wide manner. We asked whether Stau1 knockdown also affects the viral mimicry state induced by decitabine. First, we examined the effect of Stau1 knockdown on the activation status of PKR and its downstream signaling. Since ERV and other dsRNAs act as PKR-activating dsRNAs, Stau1 knockdown should decrease the phosphorylation PKR and shutdown PKR signaling even when decitabine is treated. Using immunocytochemistry, we examined PKR phosphorylation and found that in Stau1-deficient cells, pPKR and peIF2α signals were decreased dramatically (Fig. S4). The signal for total PKR and elF2α remained unchanged by Stau1 knockdown, indicating that Stau1 affects the activation status of PKR signaling by stabilizing the expression of PKR-activating dsRNAs such as ERV RNAs.

Next, we investigated the effect of Stau1 in the viral mimicry state from decitabine treatment. Using total RNA-seq on decitabine treated colorectal cancer cells, we confirmed that decitabine treatment resulted in a strong induction of many ISGs. We identified a total of 82 significantly upregulated genes that were related to immune and response to virus-related (Fig. 4A; cluster 1 and 2). In addition, gene ontology analysis of the top 300 upregulated genes using ClueGo and CluePedia revealed response to type I interferon related term (Fig. 4B). In addition to immune-related genes, decitabine treatment also increased ribosome biogenesis (cluster 3), mitochondrial gene expression (cluster 4), mitotic cell cycle (cluster 5), and mitochondrial transport (cluster 6). Moreover, decitabine treatment also significantly decreased expression of various genes, which could be clustered to protein localization to the endoplasmic reticulum (cluster 7), mRNA splicing (cluster 8), and translation (cluster 9).

**Figure 4.**
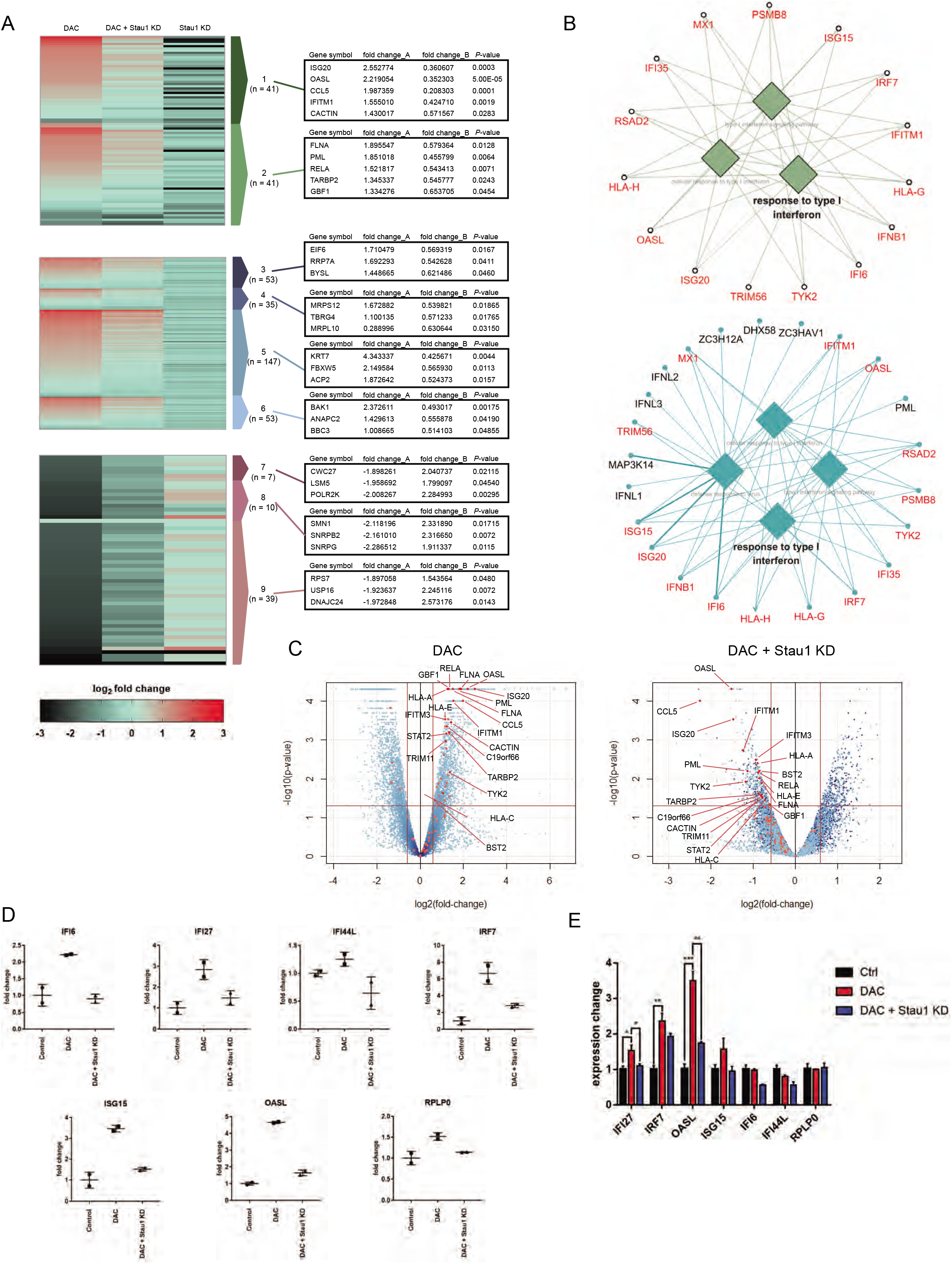
Stau1 can regulate the immune response by decitabine. (A) Heatmap of total RNA sequencing results. The first column is the RNA expression ratio of siLuc+decitabine / siLuc+DMSO, the second column is the RNA expression ratio of siStau1+decitabine / siLuc+DMSO, and the third column is the RNA expression ratio of siStau1+DMSO / siLuc+DMSO. Fold_change_A indicates the expression change of the first column while fold_change_B denotes the expression change of the second column normalized by the first column (siStau1+decitabine / siLuc+decitabine). P-values of the fold_change_B is shown in the third column. Cluster 1: related to immune, cluster 2: response to virus-related genes, cluster 3: ribosome biogenesis, cluster 4: mitochondrial gene expression, cluster 5: mitotic cell cycle, cluster 6: mitochondrial transport, cluster 7: protein localization to the endoplasmic reticulum, cluster 8: mRNA splicing, cluster 9: translation. (B) GO of top 300 upregulated genes after decitabine treatment (top) and top 500 downregulated genes in Stau1-deficient cells (bottom). Overlapping genes are marked in red. (C) Volcano plot of the sequencing data with immune response-related genes expression indicated in red. The left plot shows the RNA expression change by decitabine treatment while the right plot shows the RNA expression change in Stau1-deficient cells. (D) The RNA expression level of several IGSs in the total RNA-seq result. (E) Validation of the ISG expressions using qRT-PCR. *n*= 3 and error bars denote s.e.m. *p-value < 0.05; **p-value < 0.01 ***p-value < 0.001 ****p-value < 0.0001

We then examined the effect of Stau1 knockdown on the genes that were significantly up- or downregulated by decitabine. We found that most immune and virus-related genes in cluster 1 and 2 showed a significant decrease in the expression in Stau1-deficient cells (Fig. 4A). Indeed, the GO analysis on top 500 downregulated genes revealed response to type I interferon and other immune-related terms. This is consistent with our J2 fCLIP-seq analysis that Stau1 knockdown decreased ERV RNA expression levels and subsequently prevented the activation of immune response by decitabine. Interestingly, we found that other upregulated genes in clusters 3~6 also showed decreased expression upon Stau1 knockdown. On the contrary, genes that were downregulated by decitabine treatment were upregulated in Stau1-deficient cells (Fig. 4A). These data suggest that Stau1 knockdown virtually cancels the effect of decitabine not only in the induction of ISGs but also in other cell signaling systems.

We closely analyzed the ISGs that were regulated by decitabine and Stau1. Our volcano plot analysis reveals that most of the highly affected genes by decitabine were ISGs, and that these ISGs showed significant downregulation in Stau1-deficient cells (Fig. 4C). The fold change of the representative ISGs from the sequencing data is shown in Fig. 4D. Clearly, Stau1 knockdown eliminated the induction of ISGs by decitabine treatment. We further confirmed our high-throughput sequencing data using qRT-PCR of the same group of ISGs, which showed a similar trend for nearly all of the genes tested (Fig. 4E). Collectively, our data suggest that the Stau1 is critical in mediating the response to decitabine both in viral mimicry and others.

### Patients with low Stau1 expression show inferior response to the DNMTi therapy

Although previous and the current study mainly used colorectal cancer cell line HCT116, two DNMTis, azacitidine and its deoxyderivative decitabine, have been approved to treat patients with MDS and AML. Thus, to investigate the clinical implications of our findings, we extended our analysis to MDS and AML. We first treated low-dose decitabine in KG-1 acute myeloid leukemia cells and confirmed the induction of numerous ERV RNAs (Fig. 5A). Interestingly, although most ERV RNAs showed increased expression, the type of ERV RNAs with the highest induction was different from that of HCT116 colorectal cancer cells. Moreover, ERV9-1, which showed a strong induction in HCT116 cells, was downregulated in KG-1 cells. This indicates that it is the combined level of total cellular dsRNAs rather than the expression of a single or a group of few ERV elements that are necessary to induce the viral mimicry state by DNMTis. Under this condition, we transfected cells with an siRNA against Stau1 and examined its effects. Consistent with our earlier data, we found that the Stau1 knockdown significantly rescued cell death from decitabine in KG-1 cells as well (Fig. 5B). Moreover, the Stau1 knockdown decreased the expression of key ERV RNAs (Fig. 5C). To our surprise, the expression of ERV-L was decreased in Stau1-deficient KG-1 cells, indicating that the mode of regulation of Stau1 on individual ERV RNAs is context-dependent.

**Figure 5.**
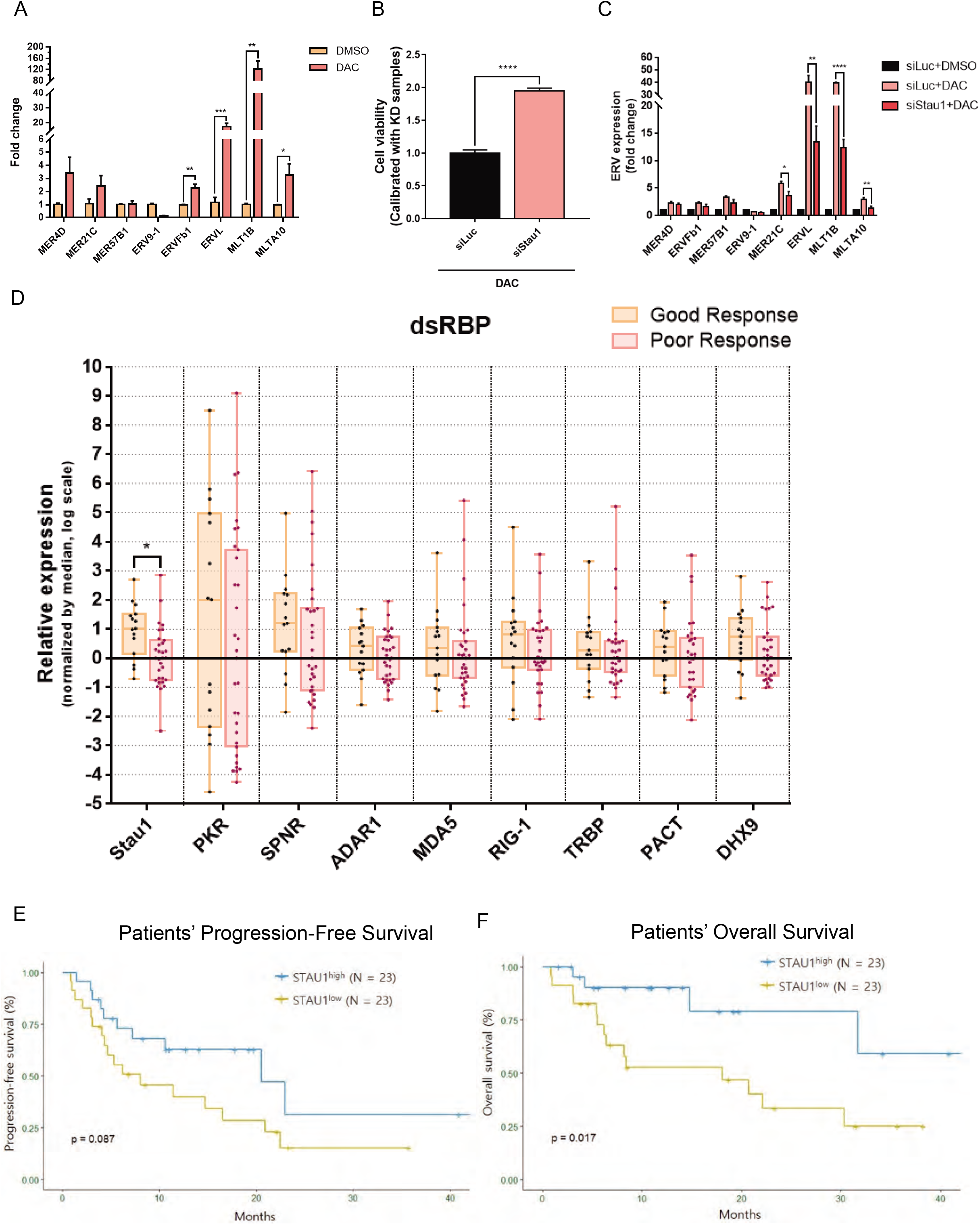
MDS and AML patients with low Stau1 level show inferior response to the DNMTi therapy. (A) Transient low dose exposure of decitabine resulted in the induction of several ERV RNAs in KG-1 AML cells. *n= 3* and error bars denote s.e.m. (B) Knockdown of Stau1 significantly rescued the cell death from decitabine treatment in KG-1 cells. n= 15 and error bars denote s.e.m. (C) Stau1 knockdown also resulted in decreased expression of several ERV RNAs examined. *n= 3* and error bars denote s.e.m. (D) Analysis of dsRBP mRNA expression from RNAs isolated from bone marrow aspirates from 46 MDS and AML patients. All patients received the DNMTi therapy, and samples were grouped based on the treatment outcome. (E, F) Patients with low Stau1 expression exhibited poor PFS (E) and OS (F) rates compared to those with high Stau1 mRNA expression. *p-value < 0.05; **p-value < 0.01 ***p-value < 0.001 ****p-value < 0.0001

We further extended our findings and analyzed samples of MDS and AML patients who underwent the DNMTi therapy. Between October 2015 and September 2018, 83 patients were diagnosed as MDS or AML and received either azacitidine or decitabine at the Seoul National University Hospital. Of these, 13 MDS patients were excluded after being found to be lower-risk MDS, and their goal of the DNMTi treatment was the correction of cytopenia rather than alternating natural disease course. Other reasons for exclusion were: lack of or poor-quality bone marrow samples (n=18) and non-evaluable to treatment response (n=6; 3 due to early death before evaluation and 3 due to follow-up loss). In the end, a total of 46 patients were analyzed. High-risk MDS comprised of 23, and the other 23 were patients with AML (Table 1).

**Table 1.** Patient baseline demographics and disease characteristics.

| Characteristic | Total<br>N (%) | MDS<br>N (%) | AML<br>N (%) |
| --- | --- | --- | --- |
| Total number of patients | 46 | 23 | 23 |
| Gender |  |  |  |
| Male | 32 (69.6) | 15 (65.2) | 17 (73.9) |
| Female | 14 (30.4) | 8 (34.8) | 6 (26.1) |
| Median age, years (range) | 70 (35-89) | 67 (35-84) | 73 (65-89) |
| Disease |  |  |  |
| MDS | 23 (50.0) |  |  |
| AML | 23 (50.0) |  |  |
| De novo AML | 18 (39.1) |  |  |
| Secondary AML | 5 (10.9) |  |  |
| ECOG performance status |  |  |  |
| 1 | 19 (41.3) | 9 (39.1) | 10 (43.5) |
| 2 | 18 (39.1) | 8 (34.8) | 10 (43.5) |
| 3 | 9 (19.6) | 6 (26.1) | 3 (13.0) |
| Hypomethylating agent |  |  |  |
| Azacitidine | 11 (23.9) | 10 (43.5) | 1 (4.3) |
| Decitabine | 35 (76.1) | 13 (56.5) | 22 (95.7) |
| MDS IPSS risk |  |  |  |
| Intermediate-2 |  | 17 (73.9) |  |
| High |  | 6 (26.1) |  |
| AML European LeukemiaNet risk |  |  |  |
| Favorable |  |  | 3 (13.0) |
| Intermediate |  |  | 15 (65.2) |
| Adverse |  |  | 5 (21.7) |
| Cell counts |  |  |  |
| Hemoglobin (range) | 8.3 (2.5-11.5) | 7.6 (2.5-11.4) | 8.6 (5.6-11.5) |
| White blood cell (range) | 3045 (550-143590) | 2950 (1160-21000) | 3460 (550-143590) |
| Absolute neutrophil count (range) | 811 (3-7520) | 600 (3-5859) | 1200 (23-7520) |
| Platelet (range) | 57.5 (4-564) | 60 (4-564) | 54 (7-146) |
| BM blast (range) | 18.1 (0.8-97.4) | 8.8 (0.8-15.7) | 5 (20.4-97.4) |
| Karyotype |  |  |  |
| Complex | 14 (30.4) | 10 (43.5) | 4 (17.4) |
| Monosomal | 11 (23.9) | 9 (39.1) | 2 (8.7) |
| RUNX1 mutation |  |  |  |
| Yes | 8 | 6 | 2 |
| No | 23 | 10 | 13 |
| ASXL1 mutation |  |  |  |
| Yes | 7 | 5 | 2 |
| No | 24 | 11 | 13 |
| TP53 mutation |  |  |  |
| Yes | 10 | 8 | 2 |
| No | 26 | 8 | 18 |
| Allogeneic hematopoietic stem cell transplantation |  |  |  |
| Yes | 9 (19.6) | 9 (39.1) | 0 (0) |
| No | 37 (80.4) | 14 (60.9) | 23 (100) |
| Objective response |  |  |  |
| Yes | 13 (28.3) | 11 (45.8) | 2 (8.7) |
| No | 33 (71.7) | 13 (54.2) | 21 (91.3) |
| Death |  |  |  |
| Yes | 18 (39.1) | 5 (21.7) | 13 (56.5) |
| No | 28 (60.9) | 18 (78.3) | 10 (43.5) |

MDS, myelodysplastic syndrome; AML, acute myeloid leukemia; ECOG, Eastern Cooperative Group;
IPSS, International Prostate Symptom Score

We extracted total RNAs from the bone marrow aspirates and analyzed the mRNA expression of several dsRBPs and ERV RNAs using qRT-PCR. First of all, we could not find any correlation between patient responses to the DNMTi therapy and ERV RNA expressions (Fig. S6A). When we examined the dsRBP mRNA expressions, we found that patients with poor response to the DNMTi therapy showed lower expression for most dsRBPs examined. Among these, Stau1 showed a statistically significant difference between the two groups (Fig. 5D). This is consistent with our image-based screening experiment that Stau1 showed the strongest rescue effects to the decitabine treatment, indicating that Stau1 is critical in decitabine-mediated cell death. When we dichotomize Stau1 expression into low (below median) versus high (equal or above median), low expression of Stau1 was associated with failure to achieve an objective response to DNMTis (Table 2). Furthermore, lower expression of Stau1 was associated with significantly inferior PFS as well as inferior OS rates (Fig. 5E and 5F).

**Table 2.**
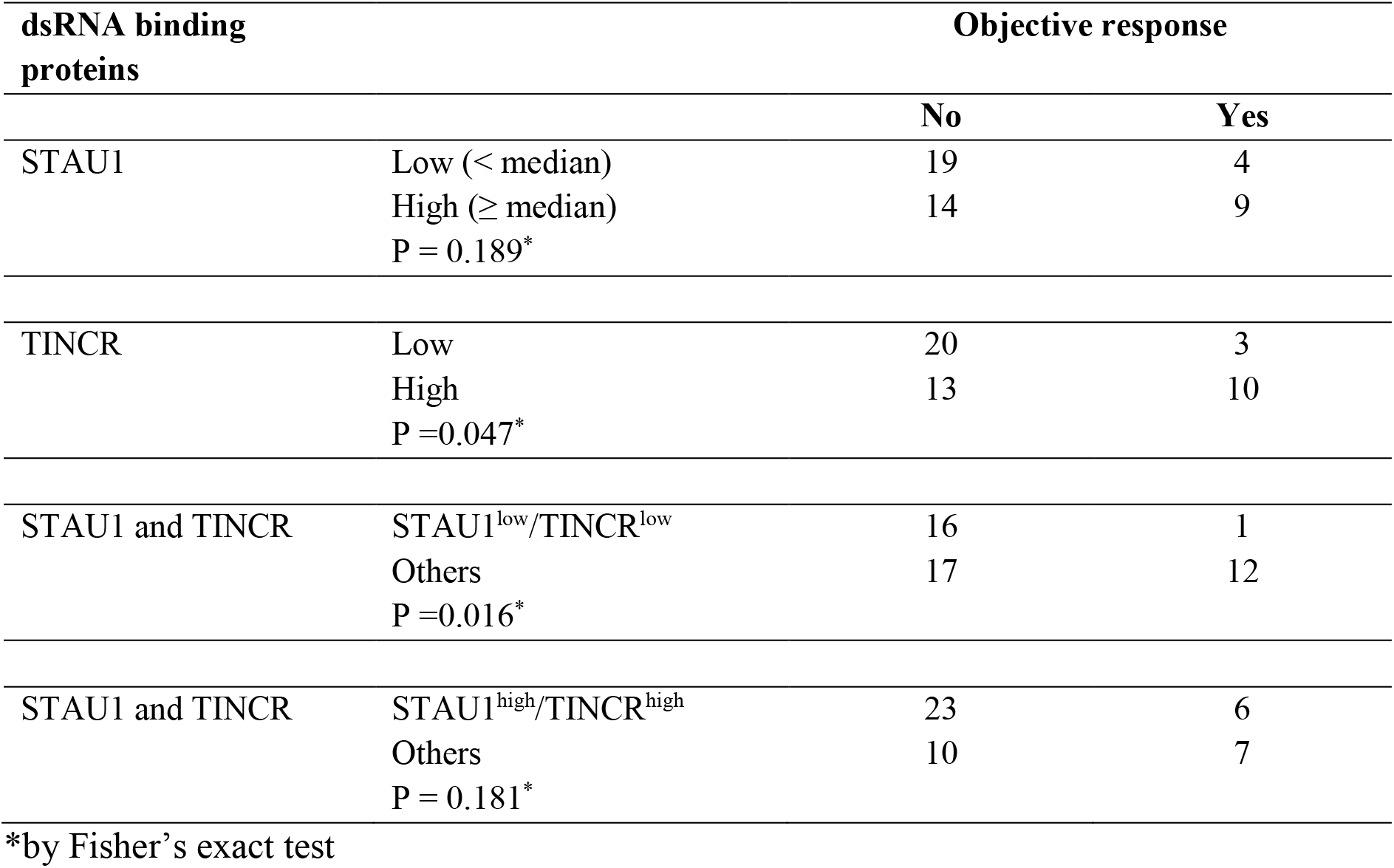
Treatment response according to dsRNA-binding protein expression and in the analyzed patients.

### Stau1 stabilizes the expression of ERV RNAs through interaction with TINCR

Lastly, we sought out the mechanism through which Stau1 stabilized ERV RNAs. Although most previous studies focused on Stau1’s binding to hairpin loops in the 3’ UTR of mRNAs to induce SMD (Park and Maquat, 2013), one study showed that Stau1 could stabilize its target mRNAs when bound with a non-coding RNA TINCR. In epidermal tissue, the TINCR-Stau1 complex mediates the stabilization of mRNAs of key genes associated with differentiation (Kretz et al., 2013). We asked whether TINCR also mediates the regulation of ERV RNAs through its interaction with Stau1. We examined TINCR RNA expression and found that TINCR RNA was expressed in HCT116 cells and was not regulated by decitabine treatment (Fig. 6A). In addition, TINCR strongly interacted with Stau1 (Fig. 6B). We then used RNAi to knockdown the TINCR expression and analyzed its effect on ERV RNA levels. We found that in TINCR-deficient cells, many of the ERV RNAs under the regulation of decitabine were downregulated (Fig. 6C). This is strikingly similar to the effects of Stau1 knockdown. We further analyzed how TINCR knockdown assisted Stau1 to stabilize ERV RNAs. We performed the Stau1 fCLIP and examined the interaction between Stau1 and ERV RNAs using qRT-PCR (Fig. 6D). We found that in TINCR-deficient cells, the interaction between Stau1 and ERV RNAs was significantly decreased (Fig. 6E). This indicates that TINCR enhances the binding between Stau1 and ERV RNAs, which may allow Stau1 to protect ERV RNAs from degradation.

**Figure 6.**
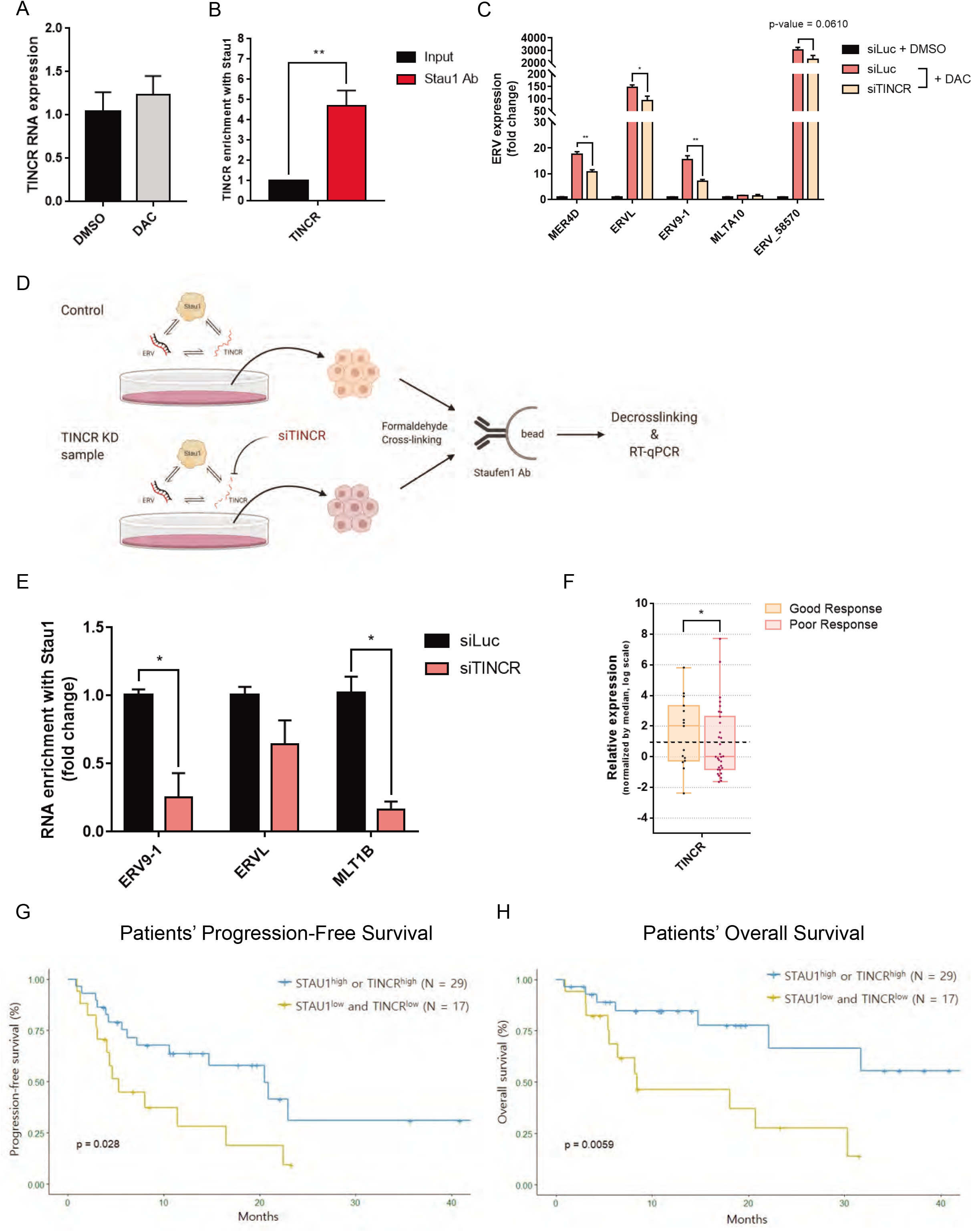
TINCR non-coding RNA assists Stau1 in stabilizing ERV RNAs. (A) TINCR RNA expression in HCT116 cells was analyzed through qRT-PCR. *n*= 3 and error bars denote s.e.m. (B) Stau1 RNA-IP after formaldehyde crosslinking revealed that Stau1 strongly interacted with TINCR. *n*= 3 and error bars denote s.e.m. (C) Knockdown of TINCR using an siRNA resulted in decreased expression of ERV RNAs. *n*= 3 and error bars denote s.e.m. (D) A schematic representation of the Stau1 fCLIP process after knockdown of TINCR using an siRNA. (E) In TINCR-deficient cells, the interaction between Stau1 and ERV RNAs was decreased. *n= 3* and error bars denote s.e.m. (F) Analysis of TINCR RNA expression in 46 MDS/AML patient samples. Patients with poor response to the DNMTi therapy showed lower expression of TINCR. (G, H) Patients with low Stau1 and TINCR expressions exhibited inferior PFS (G) and OS (H) rates compared to those with high Stau1 or TINCR RNA expressions.

To investigate the clinical significance of TINCR expression, we measured the TINCR RNA level in the bone marrow aspirates from 46 patients who underwent the DNMTi therapy. Similar to that of Stau1, TINCR expression was low in patients with poor response to DNMTis (Fig. 6F). Moreover, lower expression of Stau1 and TINCR was associated with significantly inferior PFS and OS (Fig. 6G and 6H) compared to their counterparts. In multivariate analyses, Stau1^low^ TINCR^low^ was significantly associated with reduced PFS (hazard ratio [HR] 2.72, 95% confidence interval [CI] 1.23-6.01; p-value = 0.014) and OS (HR 4.97, 95% CI 1.73-14.27; p-value = 0.003), respectively (Table 3 and 4). Stau1^low^ TINCR^low^ group also maintained the prognostic significance both in patients with or without complex karyotype in PFS and OS (Fig. S6B-S6D).

**Table 3.** Variables associated with progression free survival in patients with higher-risk myelodysplastic syndrome or acute myeloid leukemia.

| Variables | Univariate analysis |  | Multivariate analysis |  |
| --- | --- | --- | --- | --- |
|  | HR (95% CI) | *p* * | HR (95% CI) | *p* * |
| STAU1<sup>low</sup>/TINCR<sup>low</sup> | 23.0 (1.07-4.92) | 0.033 | 2.72 (1.23-6.01) | 0.014 |
| Age (<70 vs ≥70) | 0.74 (0.33-1.64) | 0.458 |  |  |
| Sex (Male vs Female) | 1.79 (0.77-4.15) | 0.176 |  |  |
| Disease (MDS vs AML) | 0.85 (0.38-1.91) | 0.697 |  |  |
| ECOG PS (0-1 vs 2-3) | 2.70 (1.14-6.42) | 0.025 | 2.41 (1.00-5.81) | 0.050 |
| HMA (AZA vs DAC) | 0.96 (0.38-2.41) | 0.932 |  |  |
| Complex karyotype | 2.60 (1.11-6.10) | 0.029 | 2.95 (1.21-7.23) | 0.018 |
| Monosomal | 2.25 (0.94-5.42) | 0.070 |  |  |
| RUNX1 mutation | 1.02 (0.37-2.84) | 0.969 |  |  |
| ASXL1 mutation | 1.05 (0.35-3.16) | 0.930 |  |  |
| TP53 mutation | 1.15 (0.44-3.01) | 0.775 |  |  |

HR, hazard ratio; CI, confidence interval; MDS, myelodysplastic syndrome; AML, acute myeloid leukemia; ECOG PS, Eastern Cooperative Group Performance Status; AZA, Azacitidine; DAC, Decitabine
\*P values were derived from the Cox proportional regression model including all variables in the table.

**Table 4.** Variables associated with allogeneic hematopoietic stem cell transplantation-censored overall survival in patients with higher-risk myelodysplastic syndrome or acute myeloid leukemia.

| Variables | Univariate analysis |  | Multivariate analysis |  |
| --- | --- | --- | --- | --- |
|  | HR (95% CI) | *P* * | HR (95% CI) | *P* * |
| STAU1<sup>low</sup>/TINCR<sup>low</sup> | 3.70 (1.36-10.04) | 0.010 | 4.97 (1.73-14.27) | 0.003 |
| Age (<70 vs ≥70) | 1.63 (0.53-5.00) | 0.393 |  |  |
| Sex (Male vs Female) | 1.93 (0.70-5.33) | 0.203 |  |  |
| Disease (MDS vs AML) | 1.76 (0.62-5.05) | 0.289 |  |  |
| ECOG PS (0-1 vs 2-3) | 4.36 (1.26-15.08) | 0.020 | 3.15 (0.88-11.26) | 0.077 |
| HMA (AZA vs DAC) | 1.14 (0.37-3.48) | 0.815 |  |  |
| Complex karyotype | 4.45 (1.50-13.21) | 0.007 | 5.82 (1.80-18.87) | 0.003 |
| Monosomal | 3.01 (1.03-8.81) | 0.045 |  |  |
| RUNX1 mutation | 0.50 (0.11-2.25) | 0.364 |  |  |
| ASXL1 mutation | 0.81 (0.18-3.66) | 0.779 |  |  |
| TP53 mutation | 2.16 (0.68-6.87) | 0.192 |  |  |

HR, hazard ratio; CI, confidence interval; MDS, myelodysplastic syndrome; AML, acute myeloid leukemia; ECOG PS, Eastern Cooperative Group Performance Status; AZA, Azacitidine; DAC, Decitabine
\*P values were derived from the Cox proportional regression model including all variables in the table.

### Staul-TINCR mediates the efficacy of decitabine by stabilizing ERV RNAs

Taken together, our study establishes Stau1 as a key regulatory factor whose expression is critical in the effective DNMTi therapy. In our model, the transcription of ERV RNAs is increased by exposure to decitabine through the demethylation of their promoters. When transcribed, these RNAs are recognized by the Stau1-TINCR complex, which affects their subcellular localization (in the case of ERV-L in HCT116 cells) or stability. The accumulation of stabilized ERV RNAs can then trigger the downstream immune response mediated by PKR and MDA5, which induce interferons and transform the cells into the viral mimicry state (Fig. 7). Therefore, Stau1 can regulate an immune response by cellular dsRNAs by regulating their stability and subcellular localization.

**Figure 7.**
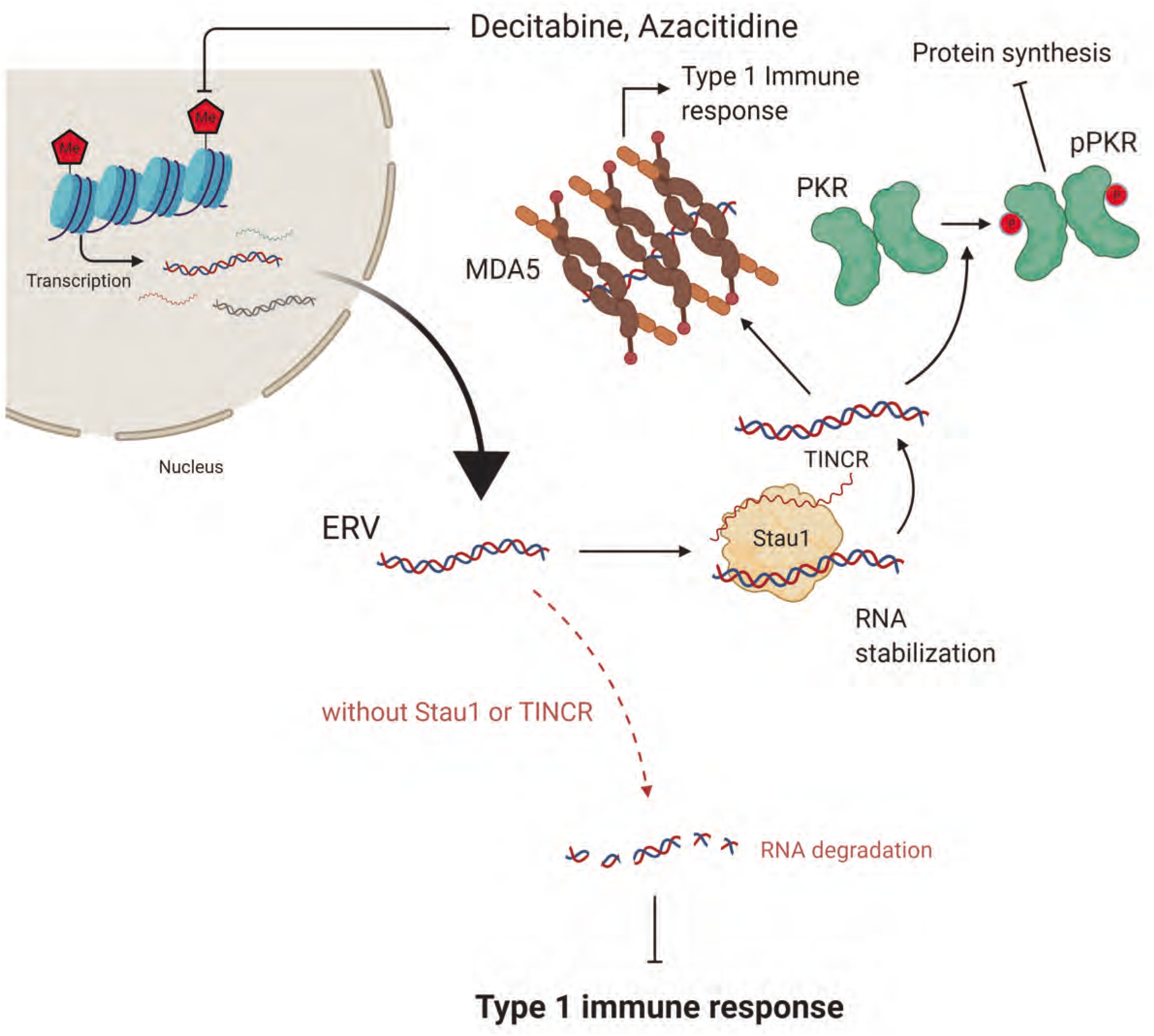
Stau1 mediates the immune response by the DNMTi treatment by stabilizing ERV RNAs. Demethylation due to DNMTi treatment results in the induction of many cellular dsRNAs, including ERV RNAs. These RNAs are stabilized by Stau1 in complex with TINCR, which leads to the activation of dsRNA sensors of innate immune response proteins such as MDA5 and PKR. In the absence of Stau1 or TINCR, ERV RNAs are destabilized, which fails to trigger an immune response, making the therapy ineffective.

## DISCUSSION

Our present data provide a non-canonical role of Stau1 in regulating the immune response to the DNMTi therapy by stabilizing ERV RNAs. Previous studies have established that DNMTis reduced the proliferation of cancer-initiating cells by activating dsRNA sensors of the innate immune system, converting the cells into the viral mimicry state (Chiappinelli et al., 2015; Roulois et al., 2015). However, it remains unclear how these dsRNAs are being regulated at the post-transcriptional level. Our data reveal that several dsRBPs may recognize ERV RNAs whose expressions were suppressed by DNA methylation in normal state. Remarkably, we found that the stability and/or subcellular localization of these immune triggering dsRNAs were under the regulation of Stau1 and TINCR. More importantly, AML and MDS patients with lower expression of Stau1 and TINCR showed significantly inferior response to the DNMTi therapy, suggesting that understanding the regulation of cellular dsRNAs by RNA-binding proteins may provide the foundation for developing a patient specific treatment strategy.

Both HCT116 colorectal cancer cells and KG-1 AML cells showed good sensitivity to the transient low dose exposure to decitabine. In both cells, decitabine treatment resulted in strong induction of ERV RNAs and subsequent activation of viral defense response. Interestingly, however, the gene identity of ERV elements that were regulated by decitabine was clearly different. Under our experimental conditions, ERV9-1 and ERV-L were most strongly induced in HCT116 cells while in KG-1 cells, ERV-L and MLT1B were induced by more than 30 folds. These results also indicate that ERV RNAs responsive to the DNMTi treatment may differ from patient to patient. Indeed, previous clinical and fundamental studies could not pin-point the ERV RNA that could be used as a response marker for the DNMTi therapy. Considering that most dsRBPs, including the dsRNA sensors of the innate immune system, recognize the length rather than specific sequences of the target RNA, the relevant marker might be the collective expression of all long dsRNAs. Previously, we developed an RNA-intercalator based sensor system that could detect total dsRNA expression in cells (Ali et al., 2020). Indeed, the expression of total dsRNAs isolated from HCT116 cells treated with decitabine showed an increasing trend over time (Ali et al., 2019). Together with dsRBP expression, profiling the dynamic induction of total dsRNAs may provide valuable insights toward developing clinical biomarkers to predict the possible outcome of the DNMTi therapy.

Although many studies on decitabine and azacitidine have been focused on ERV RNAs, our J2 fCLIP-seq analysis clearly showed that other cellular dsRNAs were also involved. Notably, the fraction of ERV RNAs in our sequencing reads in the decitabine treated sample was decreased compared to that of the control sample, despite the strong induction of ERV RNAs. This indicates that other cellular dsRNAs were also regulated, and most likely to a greater degree, by decitabine. We have previously shown that inverted Alu repeats occupy the largest fraction of PKR-activating cellular dsRNAs in human cells (Kim et al., 2018). In addition, LINE RNAs can also form doublestranded secondary structures that can activate PKR and trigger the immune response (Kim et al., 2019; Kim et al., 2018). Our J2-seq data clearly show that both Alu repeats, as well as LINE and other cellular dsRNAs, are regulated by decitabine and Stau1. Of note, total RNA-seq analysis resulted in too few sequencing reads to make statistically meaningful conclusions regarding the regulation of cellular dsRNAs by decitabine and Stau1. Therefore, applying the J2-seq to examine dsRNA expression profile in various solid tumors and contexts may provide a better experimental approach in unraveling the molecular effects of the DNMTi treatment.

DNMTis are receiving increasing attention to be used as a chemotherapy drug to treat solid tumors in addition to MDS and AML. Recent studies have shown that DNMTis and the subsequent immune response can potentiate sensitivity to immune checkpoint blockade and other immunotherapies (Chiappinelli et al., 2015; Roulois et al., 2015; Wang et al., 2019). However, the identification of responsible dsRNAs and investigation on the post-transcriptional regulation of these dsRNAs should be the precedent. Although our study is focused on Stau1 and TINCR, our dsRBP screening data clearly reveal that multiple dsRBPs such as Stau1 and SPNR may work together or work against each other such as in the case of Stau1 and PACT or DHX9. Moreover, other yet to be identified, dsRBPs may also regulate the cellular response to DNMTis. Rigorous investigation on the dsRBP-dsRNA regulatory network and the validation of the translational connotations through large clinical cohorts will greatly expand the applicability of DNMTis and cellular dsRNAs in fighting cancer.

## ACKNOWLEDGEMENTS

We thank the members of our laboratories for their input and helpful discussions. This research was supported by the Korean government Ministry of Science and ICT (NRF-2019R1C1C1006672 to Y.K.) (NRF-2013M3A9B5076486 to D.S.) and the KAIST Future Systems Healthcare Project (KAISTHEALTHCARE42 to Y.K.).

## AUTHORS CONTRIBUTIONS

Y.K. (Yongsuk)., J.-H.P., R.C., J.H., and Y.K(Yoosik). designed experiments. Y.K(Yongsuk)., R.C., Y.L., and M.K. performed experiments. K.H. assisted in the generation of the KO cell line. S.Y.B. and D.S. performed the RNAi screening experiment. Y.K(Yongsuk)., R.C., and H.-M.P. performed bioinformatics analyses. J.-H.P., Y.L., D.-Y.S., Y.K(Youngil)., S.-S.Y, and J.H. designed and analyzed patient samples. Y.K(Yongsuk)., J.-H.P., R.C., J.H., and Y.K(Yoosik). wrote the manuscript. All authors analyzed the data.

## DECLARATION OF INTERESTS

The authors declare that they have no competing interests.

## STAR*METHODS

### RESOURCE AVAILABILITY

#### Lead Contact

Further information and requests for resources and reagents should be directed to and will be fulfilled by the Lead Contact, Yoosik Kim

#### Materials Availability

All unique/stable reagents generated in this study are available from the Lead Contact with a completed Materials Transfer Agreement.

#### Data and Code Availability

The accession number for the high-throughput sequencing data reported in this paper is NCBI GEO DataSets: GSE152258.

### EXPERIMENTAL MODEL AND SUBJECT DETAILS

#### Cell lines

Wild-type HCT116 and Staufen1 knockout (KO) HCT116 cells were cultured in RPMI media (Welgene) supplemented with 10% Fetalgro bovine growth serum (RMbio). KG-1 cells were grown in RPMI-1640 (Welgene) supplemented with 10% fetal-bovine serum (Thermo Fisher). All cells were incubated at 37 °C in a humidified atmosphere of 5 % CO2. All cell lines used in this study were male. All cell lines were authenticated using short tandem repeat (STR) profiling(PowerPlex 1.2; Promega), and results were compared with reference STR profiles available through the ATCC.

#### Human bone marrow aspirates

To investigate whether pretreatment dsRBP expression is associated with treatment outcomes in patients with newly diagnosed higher-risk MDS (HR-MDS; Intermediate-2 or High-risk group according to the International Prognostic Scoring System [IPSS] (Greenberg et al., 1997) or AML and received DNMTis as the frontline therapy, RNAs were extracted from the patients’ bone marrow aspirates that were cryopreserved with RNAlater stabilizer before initiation of the DNMTi treatment. Subsequently, RNAs were purified using the Nucleospin RNA/Protein kit (Macherey-Nagel) following the manufacturer’s instructions. All patients were diagnosed either HR-MDS or AML according to the World Health Organization classification criteria for myeloid neoplasms (Arber et al., 2016) and received either azacitidine or decitabine. Patients who were diagnosed with lower-risk MDS or myeloid malignancies other than MDS or AML, who lacked clinical information or bone marrow samples, and who were not evaluable for response to DNMTis, were excluded from the analysis. The use of clinical data and the secondary use of preserved samples from the biorepository with all patients’ informed consent were reviewed and approved by the Institutional Review Board (IRB) of Seoul National University Hospital, Seoul, Republic of Korea (IRB number: H-1810-002-974).

### METHOD DETAILS

#### Chemical treatment

Cells were treated with 500 nM of decitabine for 24 h and incubated with fresh media for 4 additional days before analysis. For transfection, siRNAs were transfected with Lipofectamine 3000 (Thermo Fisher Scientific) following the manufacturer’s instruction. For KG-1 cells, electroporation was done in buffer R using Neon (Thermo Fisher Scientific). Electroporation was conducted in a single 20 ms pulse of 1700 V condition by following the manufacturer’s instruction. Information on siRNA sequences is provided in Table 5.

**Table 5.**
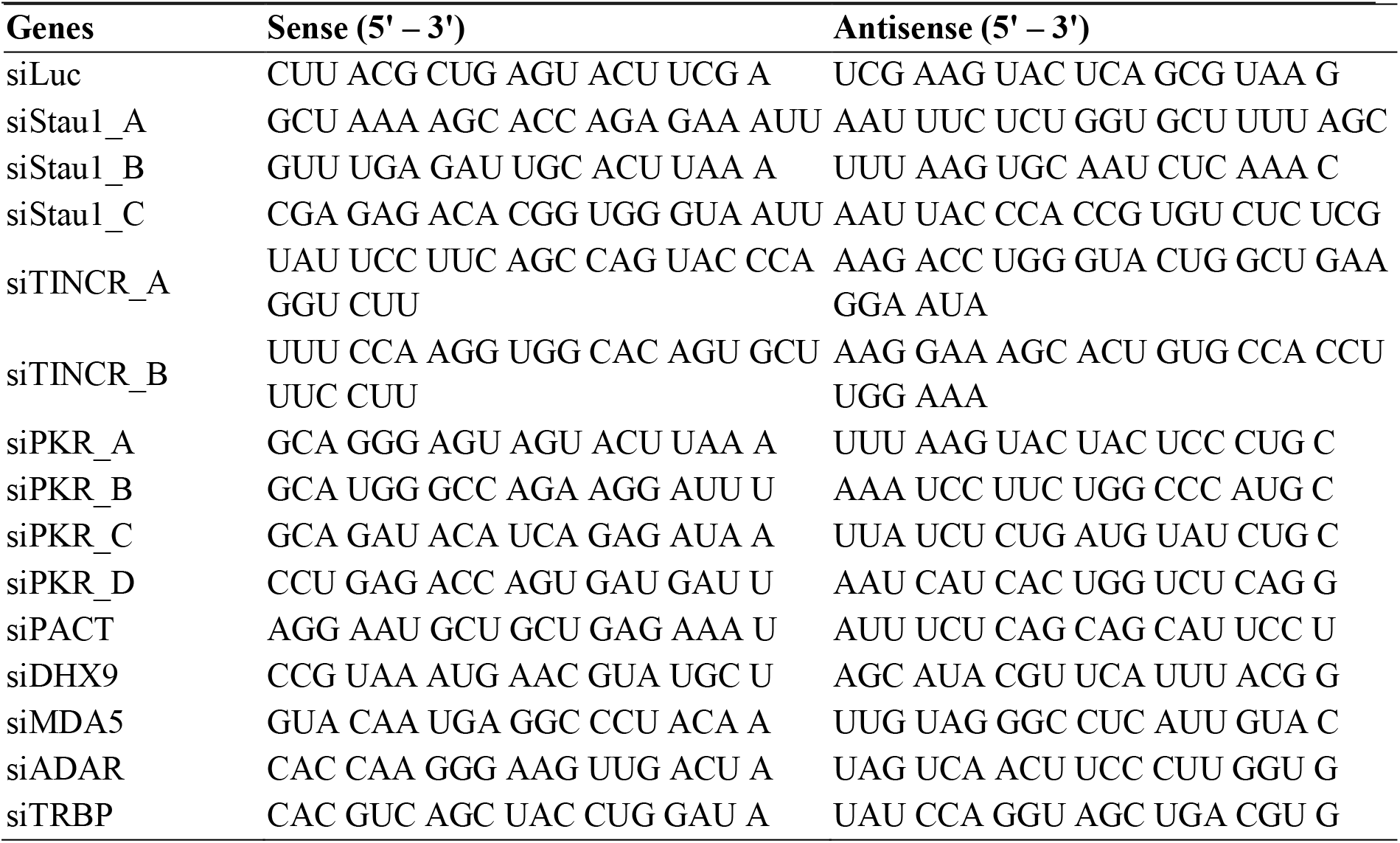
siRNA sequences.

#### RNA-FISH

RNA probes were synthesized from genomic DNA with reverse primers containing T7 promoter sequences (5’-TAATACGACTCACTATAGGG-3’). RNA was transcribed from the PCR products using MEGAscript T7 kit (Thermo Fisher Scientific) with DIG RNA labeling mix (Sigma-Aldrich). Following RNA transcription, Turbo DNase (Thermo Fisher Scientific) was added to remove the template DNA. RNA was purified using acid-phenol chloroform and hydrolyzed to ~200 bp using the hydrolysis buffer (40 mM NaHCO3 and 60 mM Na2CO3, pH 10). After incubation, stop buffer (0.2 M NaOAc, pH 6) was added to halt the hydrolysis reaction, and 200 μl of hybridization buffer (50% formamide, 10% dextran sulfate, 0.1% SDS, 300 ng/ml salmon sperm DNA, 2x SSC, and 2mM VRC) was added. Sequences for the primers used to generate RNA-FISH probes are provided in Table 6.

**Table 6.**
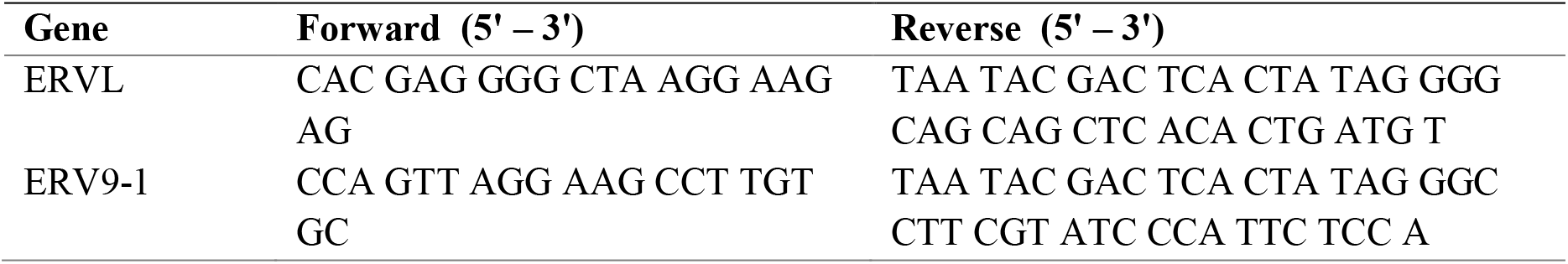
RNA-FISH probe primer sequences.

For RNA-FISH, cells were seeded on an 8-well slide and incubated for 1 day. Cells were rinsed once with phosphate-buffered saline (PBS) and fixed in 4% w/v paraformaldehyde (PFA) for 10 min at room temperature. Cells were rinsed 2 times in PBS and washed once in PBS for 5 min. To permeabilize the cells, cells were kept in 70% ethanol in 2x SSC for overnight in 4 °C. Cells were rehydrated by a successive wash in 50% ethanol and 25% ethanol for ~2-3 min each. Cells were then washed 2 times in 2x SSC for 10 min each and incubated 50% formamide for 2 h at 40 °C. Cells were hybridized with denatured probes for 4 h at 40 °C. Cells were washed 2 times in 2x SSC in 50% formamide at 37 °C and 2 times in 0.1x SSC in 50% formamide at 37 °C for 30 min each. Cells were then blocked in 0.2% BSA, 8% formamide, 2x SSC for 1 h. Sheep anti-DIG (Sigma-Aldrich) was used as the primary antibody, and fluorophore-labeled secondary antibody was used to visualize the RNAs. Cells were imaged with Carl Zeiss LSM 780 confocal microscope with a 63x objective (NA=1.40). Images were analyzed with ImageJ software.

#### Formaldehyde crosslinking and immunoprecipitation (fCLIP)

Cells were crosslinked using 0.75% PFA for 10 min at room temperature and quenched with 250 mM glycine for 10 min. Before immunoprecipitation, Immobilized Protein A Plus beads (Thermo Fisher) were washed with the wash buffer (20 mM Tris-HCl, pH 7.5, 150 mM NaCl, 10 mM EDTA, 0.1% NP-40, 0.1% SDS, 0.1% Triton X-100, 0.1% Sodium deoxycholate) and incubated with target antibody for 3 h at 4°C. Crosslinked cells were lysed in the fCLIP lysis buffer (20 mM Tris-HCl, pH, 7.5, 15 mM NaCl, 10 mM EDTA, 0.5% NP-40, 0.1% Triton X-100, 0.1% SDS, 0.1% Sodium deoxycholate) on ice for 10 min and sonicated for complete lysis. NaCl concentration was adjusted to 150 mM, and cell debris was removed through centrifugation. The lysate was immunoprecipitated by incubating with antibody-conjugated beads 3 h at 4 °C. Samples were washed 4 times with the wash buffer and eluted with the elution buffer (400 mM Tris-HCl, pH 7.5, 200 mM NaCl, 40 mM EDTA, 4% SDS, 12 M urea). Protein from the eluate was removed using proteinase K (Sigma-Aldrich) for overnight at 65°C. Eluted RNA was purified by RNA extraction. Purified RNA was either reverse transcribed and analyzed using qPCR or processed further to prepare a high-throughput sequencing library. The sequencing library was prepared and analyzed using NovaSeq by Theragen.

#### Investigations for outcomes of the DNMTi treatment

Expressions of dsRBPs were estimated by qRT-PCR and analyzed for their association with clinical characteristics and treatment outcomes. Objective treatment response to DNMTi treatment was defined as achievement of complete remission (CR), CR with incomplete hematologic recovery (CRi), or morphologic leukemia-free status according to the 2017 European LeukemiaNet (ELN) criteria (Dohner et al., 2017). As for HR-MDS, the 2017 ELN criteria was also used as this study focused on the alteration of the natural history of disease rather than hematological improvement for MDS.

Means of dsRBP RNA expression were compared by an independent two-sample t-test. Kaplan-Meier plot was used to analyze PFS from the day of DNMTi treatment initiation to the day of disease progression or death and allogeneic hematopoietic stem cell transplantation (HSCT)-censored OS from the day of DNMTi treatment initiation to the day of allogeneic HSCT or death. The prognostic effects of variables were investigated using the Cox proportional regression model.

#### RNA extraction and qRT-PCR

Total RNA was extracted using TRisure (bioline) following the manufacturer’s instruction. Purified nucleic acid was treated with DNase I (TaKaRa) to remove the genomic DNA, and RNA was reverse transcribed using RevertAid reverse transcriptase (Thermo Fisher Scientific). cDNA was analyzed using QuantStudio real-time PCR (Thermo Fisher Scientific) with SensiFAST SYBR Lo-Rox Kit (bioline). Primers used in this study are provided in Table 7.

**Table 7.**
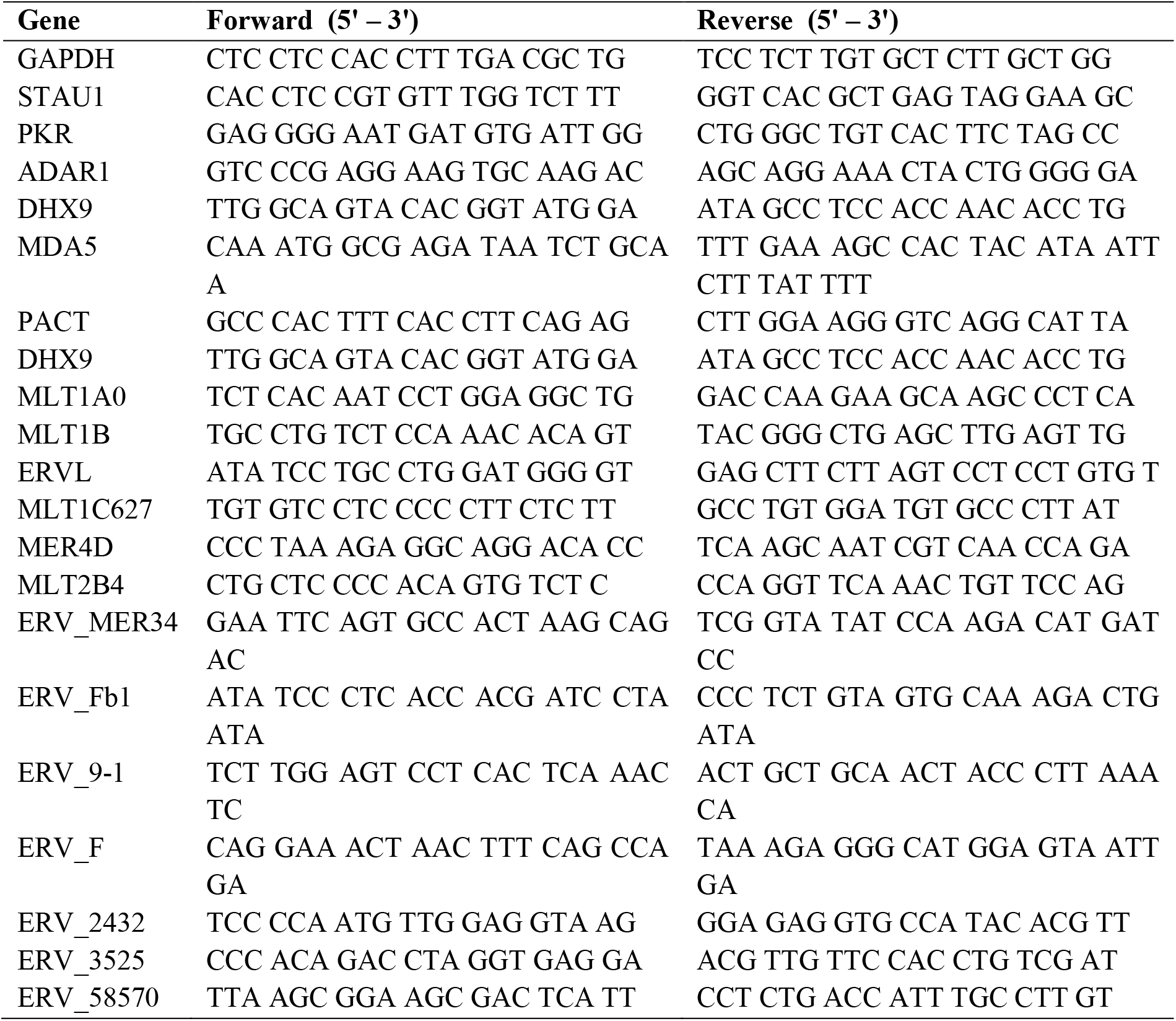
RT-qPCR primer sequences.

#### Generation of the Stau1 KO cells

Single guide RNAs for targeting human Stau1 exons were designed with a web tool, CHOPCHOP, and subcloned into a pSpCas9(BB)-2A-GFP-px458 plasmid with Cas9 protein. The KO cell line was generated following the procedure from Ran et al. (Ran et al., 2013). Briefly, a PCR product was designed with forward and reverse primers having a single guide RNA. The PCR product was inserted into the plasmid containing Cas9 protein. The plasmid was transfected to HCT116 cells using Turbofect (Thermo Fisher Scientific) following the manufacturer’s instruction. After 24 h incubation, cells with green fluorescence signals were sorted using BD FACSAria II, and single cells were seeded on a 96-well plate. Individual cells were cultured for 3 to 4 weeks, and the expression of Stau1 was examined via western blotting to isolate the KO cells.

#### Western blot

Cells were collected using a scraper and washed once with cold PBS. Cells were lysed using the RIPA buffer supplemented with protease and phosphatase inhibitors (10 mM Tris-Cl at pH 8.0, 1 mM EDTA, 1% Triton X-100, 0.1% sodium deoxycholate, 0.1% SDS, 140 mM NaCl, protease inhibitor, phosphatase inhibitor I and II, 1 mM DTT). Lysates were sonicated, and cell debris was removed through centrifugation. 30~50 μg of total protein was separated on a 10% SDS-PAGE gel and transferred to a PVDF membrane using the Amersham semidry transfer system. The following primary antibodies were used in this study: anti-PKR and anti-peIF2α were purchased from Cell Signaling Technology; anti-pPKR and anti-Stau1 were purchased from Abcam; anti-GAPDH was purchased from Santa Cruz Biotechnology.

#### Immunocytochemistry

Cells were washed once with PBS and fixed with 4% PFA for 10 min at room temperature. Fixed cells were permeabilized in 0.1% Triton X-100 and blocked in 1% BSA for 1 h. Cells were incubated in primary antibody diluted in 1% BSA for 2 h. Cells were washed 4 times with 0.1% (v/v) Tween-20 in PBS and incubated with Alex fluor conjugated secondary antibodies. Cells were imaged with a Zeiss LSM 760 confocal microscope with a C-Apochromat 63x objective (NA = 1.40). Following primary antibodies were used in this study: PKR and pPKR antibodies were purhcased from Santa Cruz Biotechnology, and eIF2α and peIF2α antibodies were from Cell Signaling Technology.

#### Sulforhodamine B (SRB) assay

Cells were fixed with 10% trichloroacetic acid solution and washed once with PBS. Cells were dried at room temperature and stained with 0.4% SRB solution (Chem Cruz). Stained cells were washed with 1% acetic acid and thoroughly dried. Cells were solubilized with 10 mM Tris (pH 10.5) and analyzed by Varioskan LUX multimode microplate reader (Thermo Fisher Scientific)

#### MTT assay

For KG-1 cells, cell viability was evaluated with MTT assay. Cells in a 96-well plate was incubated with 10 μl of 5 mg/ml MTT solution (Thermo Fisher Scientific) for 4 h. Crystals were dissolved in DMSO, and the absorbance at 590 nm was measured using Varioskan LUX multimode microplate reader (Thermo Fisher Scientific).

#### Data analysis of the J2 fCLIP-seq data

Initial transcript files were quality checked by FastaQC. Reads were aligned with the human genome (hg38) using Hisat2 (ver.2.1.0) with the cufflink option (Trapnell et al., 2012). Non-coding RNA was analyzed by counting raw transcript reads using the featurecount software (Liao et al., 2014). Each transcript was matched by a manually made ncRNA gtf reference file, which was extracted from the ncRNA database. Raw counts were normalized and analyzed with the DESeq(ver.1.36.0) software (Gentleman et al., 2004), which is capable of handling non-replicated RNA-seq data.

#### Differentially expressed gene analysis

For differential analysis, four types of treatment done to the HCT116 (siLuc DAC, siLuc DMSO, siStau1 DAC, siStau1 DMSO) were grouped and compared using both DESeq and cuffdiff (Gentleman et al., 2004; Trapnell et al., 2012). DESeq blind method, and fit-only for the dispersion of no replicate samples was used for the analysis. After the differential analysis, each RNA was sorted to one of four annotated categories; DNA, LINE, SINE, and ERV. In each category, DEG data was ordered by most significantly differentially expressed genes, most strongly upregulated, and most strongly downregulated genes for further analysis. Normalized count data of replicate RNA-seq data were also processed identically. Each DEG data of cell-line comparison was ordered by most significant, most upregulated, and most downregulated. Normalized FPKM RNA-seq data was sorted for the most significant DEG, and was compared with the previous count data for validation.

#### GO analysis of differentially expressed RNAs

Each most upregulated and downregulated RNAs was assembled into a gene list for GO analysis. KEGG pathway and GO analysis were done using the ClueGo software (Cytoscape,v.3.7.1) (Bindea et al., 2009).

### QUANTIFICATION AND STATISTICAL ANALYSIS

#### Data analysis of the RNA-seq data

Transcript files were quality checked and aligned identically. Mapped reads were assembled by two distinct methods. Initial transcript raw counts were assembled using the hg38 gtf file by the featurecount software (Liao et al., 2014). Count data with replicates were normalized and analyzed with the DESeq software (Gentleman et al., 2004). The same transcript raw counts were assembled using the cufflink software (ver.2.2.1) (Trapnell et al., 2012). Each file was converted to abundance files by cuffquant (ver.2.2.1). Abundance files were normalized by FPKM and compared by the cuffdiff (ver.2.2.1) and cuffnorm (ver.2.2.1).

#### Statistical Analysis

For statistical analyses, unpaired one-tailed Student’s t-test was used. P-value <0.05 was considered to be statistically significant. All figures show mean ± s.e.m.

## SUPPLEMENTAL FIGURE LEGENDS

**Figure S1. Decitabine treatment activates PKR signaling, related to Figure 1.** (A) A schematic for the activation of PKR signaling by decitabine. (B, C) Western blotting (B) and immunocytochemistry (C) analyses showed increased phosphorylation of PKR and its main downstream substrate eIF2α 5 days after the decitabine treatment. (D) The knockdown of PKR and MDA5 were confirmed using western blotting. (E, F) The knockdown of PKR rescued cell death from the decitabine treatment. Bright-field images (E) and SRB assay (F) clearly showed increased cell proliferation in PKR-deficient cells compared to that of the control cells 5 days after transient exposure to low dose decitabine.

**Figure S2. Decitabine treatment affects cellular dsRNA expression globally, related to Figure 2**. (A) An example of J2 fCLIP-seq read accumulation patterns mapped to an ERV locus in control, decitabine treated, and Stau1-deficient cells. ERV1_LTR1_3525 locus is shown. (B) RNAFold prediction of the secondary structure of ERV1_LTR1_3525 RNA. (C) Distribution of sequencing reads mapped to retrotransposable elements that can form dsRNAs.

**Figure S3. Generation of Stau1 KO HCT116 cells using CRISPR/Cas9 system, related to Figure 3.** A schematic of the sgRNA targeting region on the Stau1 locus. Sanger sequencing analysis shows a single nucleotide deletion, which resulted in the frameshift mutation.

**Figure S4. Stau1 affects PKR activation by decitabine, related to Figure 3.** Immunocytochemistry analysis revealed that decitabine treatment did not induce PKR and eIF2α phosphorylation in Stau1 KO cells.

**Figure S5. GO analysis of genes regulated by decitabine and Stau1, related to Figure 4.** (A) GO analysis of top 300 upregulated genes after transient low dose exposure to decitabine. (B) GO analysis of top 500 downregulated genes in Stau1-deficient cells.

**Figure S6. Analysis of MDS and AML patient samples reveal the clinical significance of Stau1 and TINCR, related to Figure 6.** (A) Examining the basal expression of several ERVs in bone marrow aspirates of MDS and AML patients before receiving the DNMTi treatment. Patients are subcategorized into good or poor group based on their response to the treatment. (B-E) Patients with low Stau1 and TINCR expressions exhibited poor PFS (B, C) and OS (D, E) regardless of complex karyotype.

## Notes

### Competing Interest Statement

The authors have declared no competing interest.

## REFERENCES

Ahmad, S., Mu, X., Yang, F., Greenwald, E., Park, J.W., Jacob, E., Zhang, C.Z., and Hur, S. (2018). Breaching Self-Tolerance to Alu Duplex RNA Underlies MDA5-Mediated Inflammation. Cell 772, 797–810 e713.

Akira, S., Uematsu, S., and Takeuchi, O. (2006). Pathogen recognition and innate immunity. Cell 724, 783–801.

Aktas, T., Avsar Ilik, I., Maticzka, D., Bhardwaj, V., Pessoa Rodrigues, C., Mittler, G., Manke, T., Backofen, R., and Akhtar, A. (2017). DHX9 suppresses RNA processing defects originating from the Alu invasion of the human genome. Nature 544, 115–119.

Ali, A.A., Kang, M., Kharbash, R., and Kim, Y. (2019). Spiropyran as a potential molecular diagnostic tool for double-stranded RNA detection. BMC Biomedical Engineering 1, 6.

Ali, A.A., Kharbash, R., and Kim, Y. (2020). Chemo- and biosensing applications of spiropyran and its derivatives - A review. Anal Chim Acta 1110, 199–223.

Arber, D.A., Orazi, A., Hasserjian, R., Thiele, J., Borowitz, M.J., Le Beau, M.M., Bloomfield, C.D., Cazzola, M., and Vardiman, J.W. (2016). The 2016 revision to the World Health Organization classification of myeloid neoplasms and acute leukemia. Blood 121, 2391–2405.

Athanasiadis, A., Rich, A., and Maas, S. (2004). Widespread A-to-I RNA editing of Alu-containing mRNAs in the human transcriptome. PLoS Biol 2, e391.

Bahn, J.H., Ahn, J., Lin, X., Zhang, Q., Lee, J.H., Civelek, M., and Xiao, X. (2015). Genomic analysis of ADAR1 binding and its involvement in multiple RNA processing pathways. Nat Commun 6, 6355.

Bindea, G., Mlecnik, B., Hackl, H., Charoentong, P., Tosolini, M., Kirilovsky, A., Fridman, W.H., Pages, F., Trajanoski, Z., and Galon, J. (2009). ClueGO: a Cytoscape plug-in to decipher functionally grouped gene ontology and pathway annotation networks. Bioinformatics 25, 1091–1093.

Chen, L.L., and Carmichael, G.G. (2008). Gene regulation by SINES and inosines: biological consequences of A-to-I editing of Alu element inverted repeats. Cell Cycle 7, 3294–3301.

Chiappinelli, K.B., Strissel, P.L., Desrichard, A., Li, H., Henke, C., Akman, B., Hein, A., Rote, N.S., Cope, L.M., Snyder, A., et al. (2015). Inhibiting DNA Methylation Causes an Interferon Response in Cancer via dsRNA Including Endogenous Retroviruses. Cell 162, 974–986.

Dhir, A., Dhir, S., Borowski, L.S., Jimenez, L., Teitell, M., Rotig, A., Crow, Y.J., Rice, G.I., Duffy, D., Tamby, C., et al. (2018). Mitochondrial double-stranded RNA triggers antiviral signalling in humans. Nature 560, 238–242.

Dohner, H., Estey, E., Grimwade, D., Amadori, S., Appelbaum, F.R., Buchner, T., Dombret, H., Ebert, B.L., Fenaux, P., Larson, R.A., et al. (2017). Diagnosis and management of AML in adults: 2017 ELN recommendations from an international expert panel. Blood 129, 424–447.

Elbarbary, R.A., Li, W., Tian, B., and Maquat, L.E. (2013). STAU1 binding 3’ UTR IRAlus complements nuclear retention to protect cells from PKR-mediated translational shutdown. Genes Dev 27, 1495–1510.

Gentleman, R.C., Carey, V.J., Bates, D.M., Bolstad, B., Dettling, M., Dudoit, S., Ellis, B., Gautier, L., Ge, Y., Gentry, J., et al. (2004). Bioconductor: open software development for computational biology and bioinformatics. Genome Biol 5, R80.

Gong, C., and Maquat, L.E. (2011). lncRNAs transactivate STAU1-mediated mRNA decay by duplexing with 3’ UTRs via Alu elements. Nature 470, 284–288.

Greenberg, P., Cox, C., LeBeau, M.M., Fenaux, P., Morel, P., Sanz, G., Sanz, M., Vallespi, T., Hamblin, T., Oscier, D., et al. (1997). International scoring system for evaluating prognosis in myelodysplastic syndromes. Blood 89, 2079–2088.

Hur, S. (2019). Double-Stranded RNA Sensors and Modulators in Innate Immunity. Annu Rev Immunol 37, 349–375.

Kim, S., Ku, Y., Ku, J., and Kim, Y. (2019). Evidence of Aberrant Immune Response by Endogenous Double-Stranded RNAs: Attack from Within. Bioessays 41, e1900023.

Kim, Y., Lee, J.H., Park, J.E., Cho, J., Yi, H., and Kim, V.N. (2014). PKR is activated by cellular dsRNAs during mitosis and acts as a mitotic regulator. Genes Dev 28, 1310–1322.

Kim, Y., Park, J., Kim, S., Kim, M., Kang, M.G., Kwak, C., Kang, M., Kim, B., Rhee, H.W., and Kim, V.N. (2018). PKR Senses Nuclear and Mitochondrial Signals by Interacting with Endogenous Double-Stranded RNAs. Mol Cell 71, 1051–1063 e1056.

Kostura, M., and Mathews, M.B. (1989). Purification and activation of the double-stranded RNA-dependent eIF-2 kinase DAI. Mol Cell Biol 9, 1576–1586.

Kretz, M., Siprashvili, Z., Chu, C., Webster, D.E., Zehnder, A., Qu, K., Lee, C.S., Flockhart, R.J., Groff, A.F., Chow, J., et al. (2013). Control of somatic tissue differentiation by the long non-coding RNA TINCR. Nature 493, 231–235.

Lander, E.S., Linton, L.M., Birren, B., Nusbaum, C., Zody, M.C., Baldwin, J., Devon, K., Dewar, K., Doyle, M., FitzHugh, W., et al. (2001). Initial sequencing and analysis of the human genome. Nature 409, 860–921.

Lemaire, P.A., Anderson, E., Lary, J., and Cole, J.L. (2008). Mechanism of PKR Activation by dsRNA. J Mol Biol 381, 351–360.

Liao, Y., Smyth, G.K., and Shi, W. (2014). featureCounts: an efficient general purpose program for assigning sequence reads to genomic features. Bioinformatics 30, 923–930.

Liddicoat, B.J., Piskol, R., Chalk, A.M., Ramaswami, G., Higuchi, M., Hartner, J.C., Li, J.B., Seeburg, P.H., and Walkley, C.R. (2015). RNA editing by ADAR1 prevents MDA5 sensing of endogenous dsRNA as nonself. Science 349, 1115–1120.

Liu, C.X., Li, X., Nan, F., Jiang, S., Gao, X., Guo, S.K., Xue, W., Cui, Y., Dong, K., Ding, H., et al. (2019). Structure and Degradation of Circular RNAs Regulate PKR Activation in Innate Immunity. Cell 177, 865–880 e821.

Masliah, G., Barraud, P., and Allain, F.H. (2013). RNA recognition by double-stranded RNA binding domains: a matter of shape and sequence. Cell Mol Life Sci 70, 1875–1895.

Park, E., and Maquat, L.E. (2013). Staufen-mediated mRNA decay. Wiley Interdiscip Rev RNA 4, 423–435.

Patel, R.C., Stanton, P., McMillan, N.M., Williams, B.R., and Sen, G.C. (1995). The interferon-inducible double-stranded RNA-activated protein kinase self-associates in vitro and in vivo. Proc Natl Acad Sci U S A 92, 8283–8287.

Pettersson, U., and Philipson, L. (1974). Synthesis of complementary RNA sequences during productive adenovirus infection. Proc Natl Acad Sci U S A 71, 4887–4891.

Ran, F.A., Hsu, P.D., Wright, J., Agarwala, V., Scott, D.A., and Zhang, F. (2013). Genome engineering using the CRISPR-Cas9 system. Nat Protoc 8, 2281–2308.

Ricci, E.P., Kucukural, A., Cenik, C., Mercier, B.C., Singh, G., Heyer, E.E., Ashar-Patel, A., Peng, L., and Moore, M.J. (2014). Staufen1 senses overall transcript secondary structure to regulate translation. Nat Struct Mol Biol 21, 26–35.

Rice, G.I., Kasher, P.R., Forte, G.M., Mannion, N.M., Greenwood, S.M., Szynkiewicz, M., Dickerson, J.E., Bhaskar, S.S., Zampini, M., Briggs, T.A., et al. (2012). Mutations in ADAR1 cause Aicardi-Goutieres syndrome associated with a type I interferon signature. Nat Genet 44, 1243–1248.

Roulois, D., Loo Yau, H., Singhania, R., Wang, Y., Danesh, A., Shen, S.Y., Han, H., Liang, G., Jones, P.A., Pugh, T.J., et al. (2015). DNA-Demethylating Agents Target Colorectal Cancer Cells by Inducing Viral Mimicry by Endogenous Transcripts. Cell 162, 961–973.

Rubin, C.M., Houck, C.M., Deininger, P.L., Friedmann, T., and Schmid, C.W. (1980). Partial nucleotide sequence of the 300-nucleotide interspersed repeated human DNA sequences. Nature 284, 372–374.

Saunders, L.R., and Barber, G.N. (2003). The dsRNA binding protein family: critical roles, diverse cellular functions. Faseb J 17, 961–983.

Schlee, M., and Hartmann, G. (2016). Discriminating self from non-self in nucleic acid sensing. Nat Rev Immunol 16, 566–580.

Thomis, D.C., and Samuel, C.E. (1993). Mechanism of interferon action: evidence for intermolecular autophosphorylation and autoactivation of the interferon-induced, RNA-dependent protein kinase PKR. J Virol 67, 7695–7700.

Trapnell, C., Roberts, A., Goff, L., Pertea, G., Kim, D., Kelley, D.R., Pimentel, H., Salzberg, S.L., Rinn, J.L., and Pachter, L. (2012). Differential gene and transcript expression analysis of RNA-seq experiments with TopHat and Cufflinks. Nat Protoc 7, 562–578.

Vukovic, L., Koh, H.R., Myong, S., and Schulten, K. (2014). Substrate recognition and specificity of double-stranded RNA binding proteins. Biochemistry 53, 3457–3466.

Wang, M., Liu, Y., Cheng, Y., Wei, Y., and Wei, X. (2019). Immune checkpoint blockade and its combination therapy with small-molecule inhibitors for cancer treatment. Biochim Biophys Acta Rev Cancer 1871, 199–224.

Weber, F., Wagner, V., Rasmussen, S.B., Hartmann, R., and Paludan, S.R. (2006). Doublestranded RNA is produced by positive-strand RNA viruses and DNA viruses but not in detectable amounts by negative-strand RNA viruses. J Virol 80, 5059–5064.

Wek, R.C., Jiang, H.Y., and Anthony, T.G. (2006). Coping with stress: eIF2 kinases and translational control. Biochem Soc Trans 34, 7–11.

Wu, B., Peisley, A., Richards, C., Yao, H., Zeng, X., Lin, C., Chu, F., Walz, T., and Hur, S. (2013). Structural basis for dsRNA recognition, filament formation, and antiviral signal activation by MDA5. Cell 152, 276–289.

