## Supplemental Figures for "Non-canonical immune response to the inhibition of DNA methylation via stabilization of endogenous retrovirus dsRNAs"

**Sup 1.**

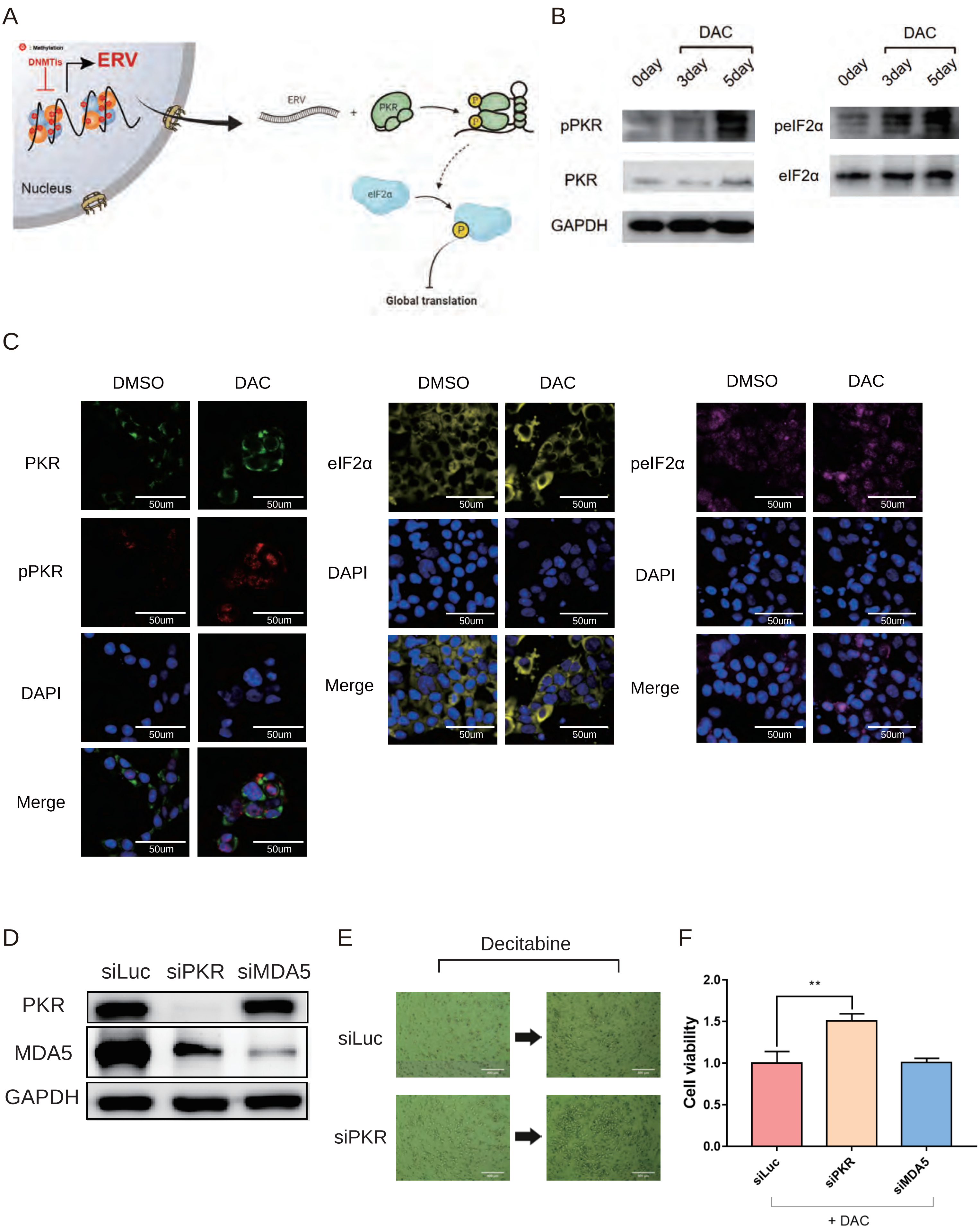
A
Methylation
DNMTis
ERV
Nucleus
ERV
PKR
P
eIF2α
P
Global translation
B
DAC
0day
3day
5day
pPKR
PKR
GAPDH
DAC
0day
3day
5day
peIF2α
eIF2α
C
DMSO
DAC
PKR
pPKR
DAPI
Merge
50um
DMSO
DAC
eIF2α
DAPI
Merge
DMSO
DAC
peIF2α
DAPI
Merge
D
siLuc
siPKR
siMDA5
PKR
MDA5
GAPDH
E
Decitabine
siLuc
siPKR
F
Cell viability
2.0
1.5
1.0
0.5
0.0
**
siLuc
siPKR
siMDA5
+ DAC

Figure S2.

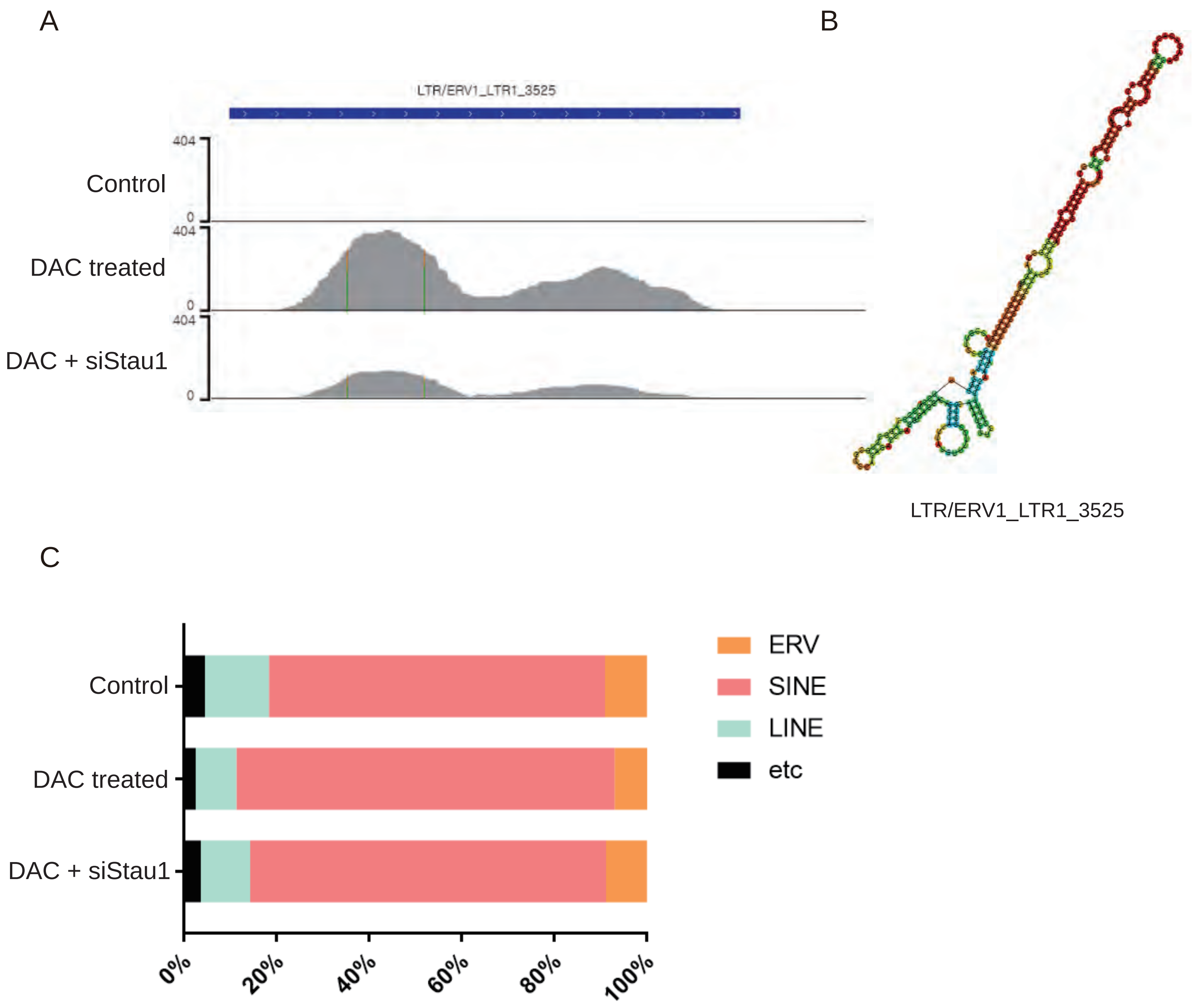
A
LTR/ERV1_LTR1_3525
404
Control
0
404
DAC treated
0
404
DAC + siStau1
0
B
LTR/ERV1_LTR1_3525
C
Control
DAC treated
DAC + siStau1
0%
20%
40%
60%
80%
100%
ERV
SINE
LINE
etc

Figure S3.

| Cell line | Sequence information |
| --- | --- |
| HCT116 Ctrl | GTGACCAAGGTTTCGGTTGGGGAGTTTGTGGGGG |
| HCT116 Stau1 KO | GTGACCAAGGTTTCGGT-GGGGAGTTTGTGGGGG |

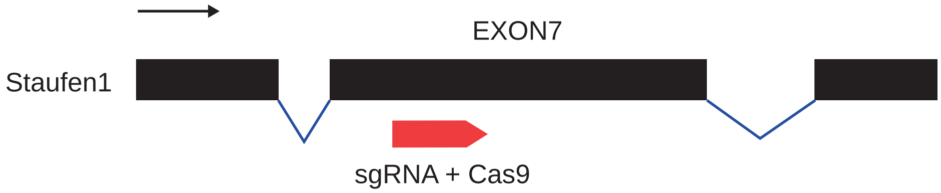

Figure S4.

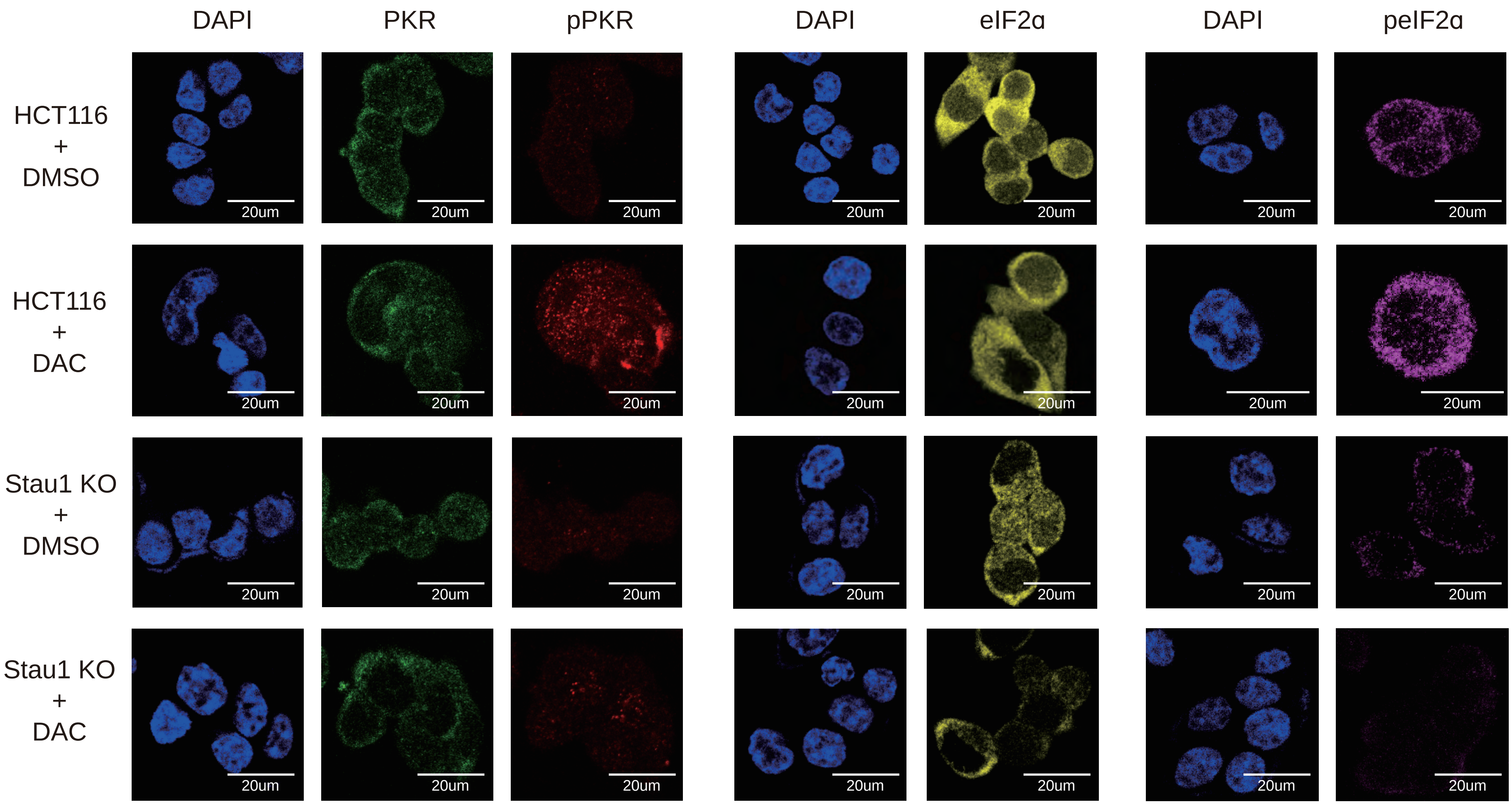
DAPI
PKR
pPKR
DAPI
eIF2α
DAPI
peIF2α
HCT116 + DMSO
HCT116 + DAC
Stau1 KO + DMSO
Stau1 KO + DAC
20um

Figure S5.

A

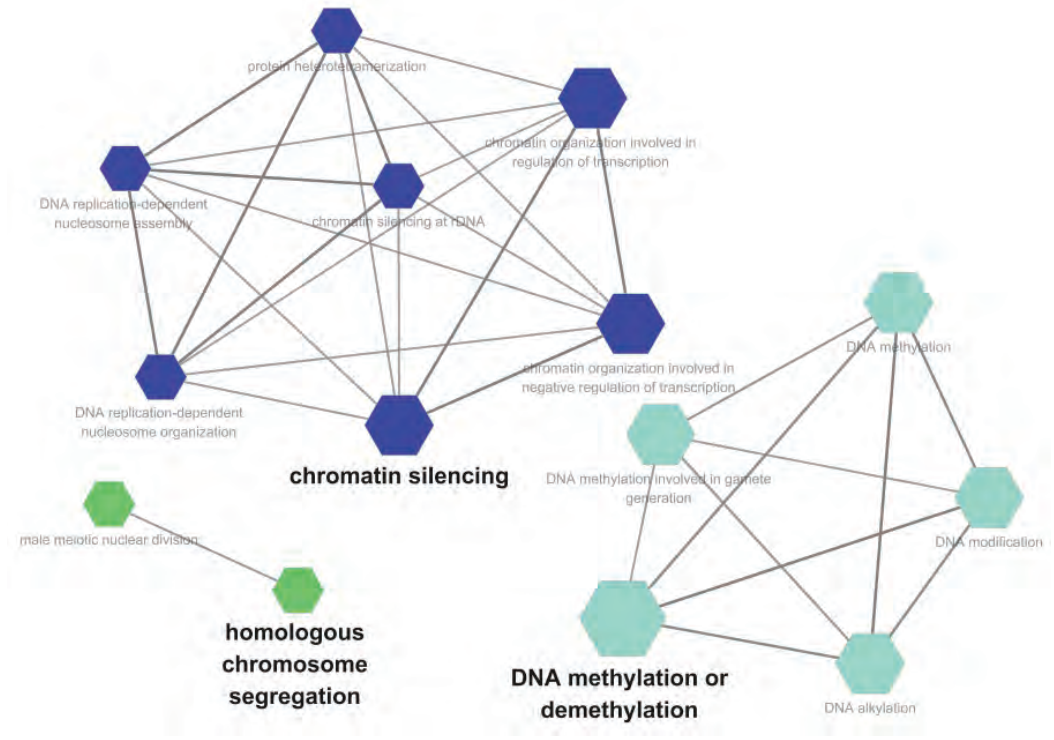
protein heterotetramerization
chromatin organization involved in regulation of transcription
DNA replication-dependent nucleosome assembly
chromatin silencing at rDNA
DNA replication-dependent nucleosome organization
chromatin organization involved in negative regulation of transcription
chromatin silencing
male meiotic nuclear division
homologous chromosome segregation
DNA methylation
DNA methylation involved in gamete generation
DNA modification
DNA methylation or demethylation
DNA alkylation

B

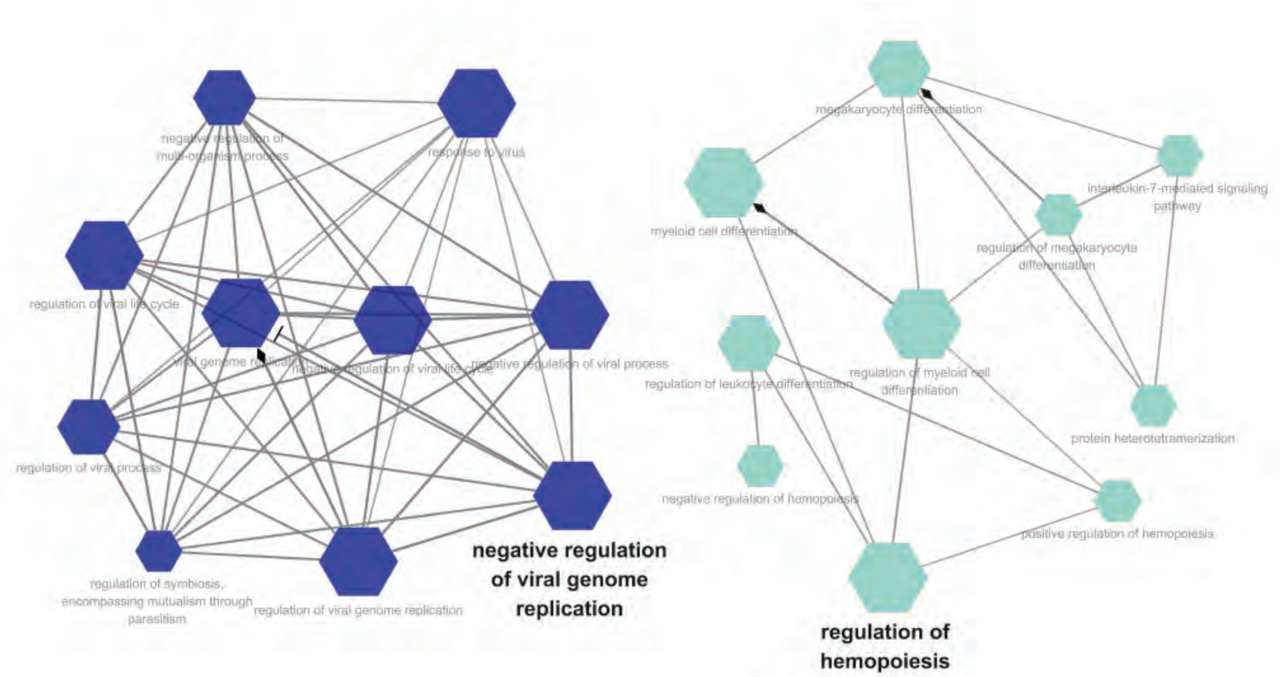
negative regulation of multi-organism process
response to virus
regulation of viral life cycle
viral genome replication
negative regulation of viral life cycle
negative regulation of viral process
regulation of viral process
regulation of symbiosis, encompassing mutualism through parasitism
regulation of viral genome replication
negative regulation of viral genome replication
megakaryocyte differentiation
myeloid cell differentiation
interleukin-7-mediated signaling pathway
regulation of megakaryocyte differentiation
regulation of leukocyte differentiation
regulation of myeloid cell differentiation
protein heterotetramerization
negative regulation of hemopoiesis
positive regulation of hemopoiesis
regulation of hemopoiesis

Figure S6.

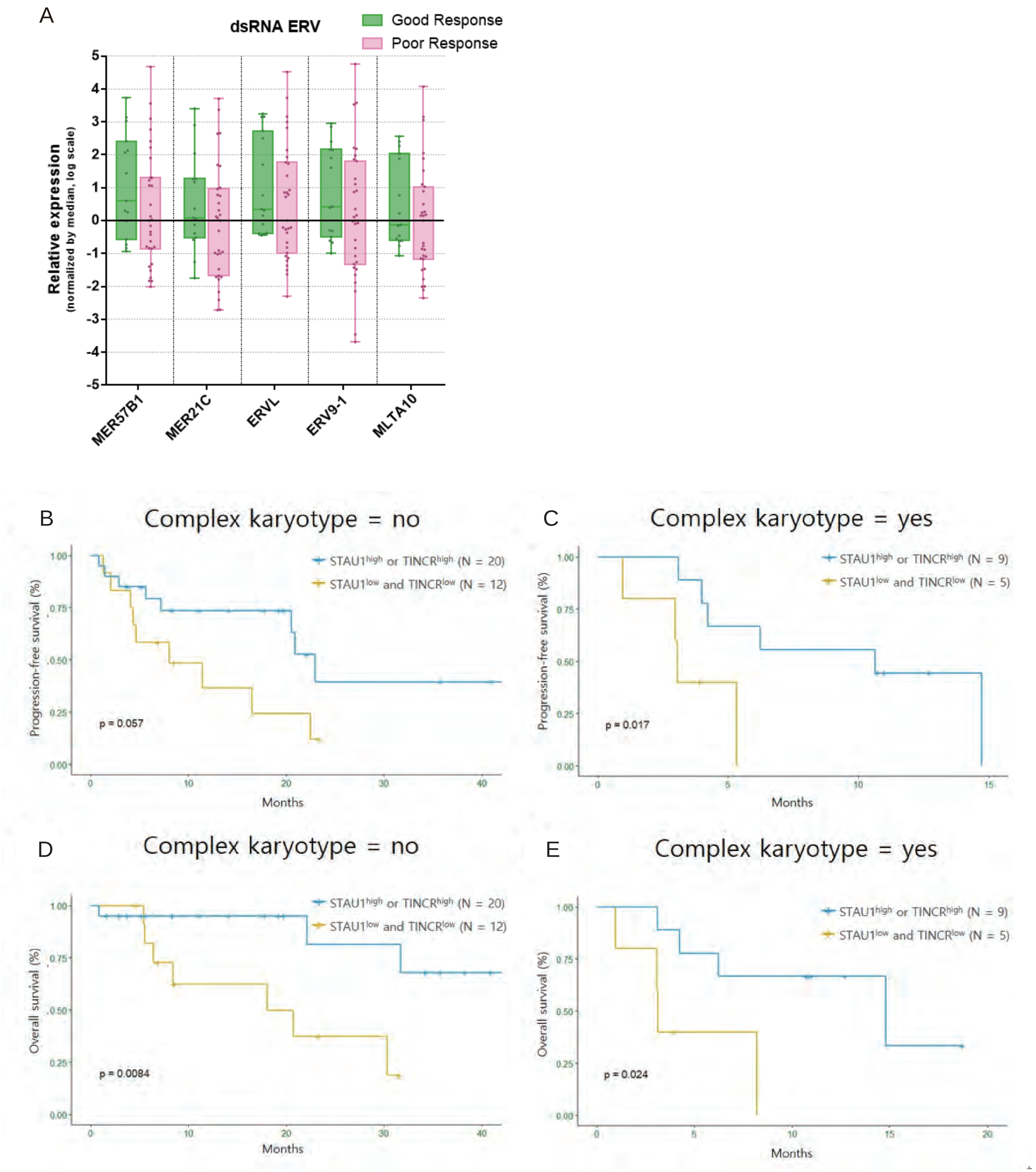
A
dsRNA ERV
Good Response
Poor Response
Relative expression
(normalized by median, log scale)
5
4
3
2
1
0
-1
-2
-3
-4
-5
MER57B1
MER21C
ERVL
ERV9-1
MLTA10
B
Complex karyotype = no
STAU1high or TINCRhigh (N = 20)
STAU1low and TINCRlow (N = 12)
Progression-free survival (%)
p = 0.057
Months
C
Complex karyotype = yes
STAU1high or TINCRhigh (N = 9)
STAU1low and TINCRlow (N = 5)
Progression-free survival (%)
p = 0.017
Months
D
Complex karyotype = no
STAU1high or TINCRhigh (N = 20)
STAU1low and TINCRlow (N = 12)
Overall survival (%)
p = 0.0084
Months
E
Complex karyotype = yes
STAU1high or TINCRhigh (N = 9)
STAU1low and TINCRlow (N = 5)
Overall survival (%)
p = 0.024
Months
